# Morphologically constrained modeling of spinous inhibition in the somato-sensory cortex

**DOI:** 10.1101/2020.08.06.230722

**Authors:** Olivier Gemin, Pablo Serna, Nora Assendorp, Matteo Fossati, Philippe Rostaing, Antoine Triller, Cécile Charrier

## Abstract

Pyramidal neurons are covered by thousands of dendritic spines receiving excitatory synaptic inputs. The ultrastructure of dendritic spines shapes signal compartmentalization but ultrastructural diversity is rarely taken into account in computational models of synaptic integration. Here, we developed a 3D correlative light-electron microscopy (3D-CLEM) approach allowing the analysis of specific populations of synapses in genetically defined neuronal types in intact brain circuits. We used it to reconstruct segments of basal dendrites of layer 2/3 pyramidal neurons of adult mouse somatosensory cortex and quantify spine ultrastructural diversity. We found that 10% of spines were dually-innervated and 38% of inhibitory synapses localized to spines. Using our morphometric data to constrain a model of synaptic signal compartmentalization, we assessed the impact of spinous versus dendritic shaft inhibition. Our results indicate that spinous inhibition is locally more efficient than shaft inhibition and that it can decouple voltage and calcium signaling, potentially impacting synaptic plasticity.

## INTRODUCTION

In the mammalian cortex, the vast majority of excitatory synapses are formed on dendritic spines, small membrane protrusions that decorate the dendrites of pyramidal neurons (PNs) [1–3]. Dendritic spines are composed of a bulbous head connected to the dendritic shaft by a narrow neck [4,5]. They exist in a large variety of shapes and sizes along individual dendrites. Spine head volume can vary between 3 orders of magnitude (0.01-1.5 μm^3^), neck length between 0.2 μm and 3 μm, and minimal neck diameter between 20 and 500 nm [6]. Spine heads are typically contacted by an excitatory synaptic input and harbor an excitatory postsynaptic density (ePSD) that contains glutamatergic α-amino-3-hydroxy-5-methyl-4-isoxazolepropionic acid (AMPA) and N-Methyl-D-aspartate (NMDA) neurotransmitter receptors, scaffolding proteins, adhesion molecules and a complex machinery of proteins undertaking the transduction of synaptic signals. The size of the spine head correlates with the size of the ePSD and the strength of synaptic transmission [7–11]. In addition to the ePSD, spines contain ribosomes, which mediate local protein synthesis, and endosomes, which play a critical role in membrane and receptor trafficking [12,13]. The largest spines often contain a spine apparatus (SA), which contributes to calcium signaling and synaptic plasticity [12,14], and some spines, especially in the upper layers of the cortex, also house an inhibitory postsynaptic specialization [15]. Spine necks are diffusional barriers that biochemically isolate spine heads from their parent dendrite [16–19]. In addition, they can filter the electrical component of synaptic signals and amplify spine head depolarization [20–22] (but see [23–25]). Both spine heads and spine necks are remodeled depending on neuronal activity [9,26,27] and in pathology [28,29]. While the relationship between spine morphology and spine function is widely acknowledged, and although dendritic spines are known to participate in different neural circuits depending on their location in the dendritic tree [30], the extent of synaptic ultrastructural diversity along individual identified dendrites has not been quantified, and the consequences of this variability on signal compartmentalization and dendritic integration remain to be investigated.

Dendritic signaling can be modeled based on anatomical and biophysical parameters [31] using “realistic” multi-compartment models [32]. These models were pioneered by Wilfrid Rall following the seminal works of Hodgkin and Huxley [33,34]. They have provided a powerful theoretical framework for understanding dendritic integration [35], spine function [36], inhibitory signaling [37,38] and electrical compartmentalization in spines [22,39,40]. However, spines and synapses are usually modeled with *ad hoc* or averaged biophysical parameters, which limits the accuracy of the prediction [41]. Modeling the actual behavior of dendritic spines requires an accurate description of their ultrastructural heterogeneity with a cell type and dendritic type resolution. To acquire such data, it is necessary to combine the nanometer resolution of electron microscopy (EM) with an approach that allows the identification of the origin of dendritic spines (i.e. location on the dendrite, type of dendrite and type of neuron) without obscuring the intracellular content. This task is arduous: 1 mm^3^ of mouse cortex contains over 50,000 of neurons, each of which establishes approximately 8,000 synaptic connections with neighboring neurons, and these synapses are highly specific, connecting multiple neuronal subtypes from various brain regions [42–45]. Reconstructing selected dendritic spines and synaptic contacts along dendritic trees requires either enormous volumes of 3D-EM acquisitions using resource-consuming approaches adapted from connectomics [46–49], or combining EM with a lower-scale imaging modality, such as confocal or 2-photon light microscopy (LM), to guide 3D-EM image acquisitions to the region of interest (ROI) [50–52]. While very powerful *in vitro* [50,53,54], correlative light-electron microscopy (CLEM) is difficult to implement in brain tissues [55–57]. New protocols are required to facilitate the *in situ* identification of targeted dendrites and synapses in different imaging modalities and to make 3D-CLEM more accessible to the neuroscientific community.

Here, we have developed a CLEM workflow combining confocal light microscopy with serial block-face scanning EM (SBEM) and targeted photo-precipitation of 3,3-diaminobenzidine (DAB) to facilitate ROI recovery. We applied this workflow to reconstruct dendritic spines located exclusively on the basal dendrites of genetically-labelled PNs in layers 2/3 (L2/3) of the somatosensory cortex (SSC) of adult mice. We analyzed the variability of their ultrastructure and estimated the electrical resistance of their neck. We also examined the distribution and the morphology of inhibitory synapses. We specifically examined dendritic spines receiving both excitatory and inhibitory inputs, which represented 10% of all spines along basal dendrites. These dually-innervated spines (DiSs) exhibited wider heads and larger ePSDs than singly-innervated spines (SiSs), and they were more electrically isolated from the dendritic shaft than SiSs of comparable head size. We then used our measurements to constrain a multi-compartment model of synaptic signaling and compartmentalization in dendrites. We assessed the effects of individual excitatory and inhibitory signals on membrane voltage and calcium concentration depending on inhibitory synapse placement (i.e. on a spine head or on the dendritic shaft) and input timing. Our results challenge the view that spinous inhibition strictly vetoes single excitatory inputs and rather suggest that it fine-tunes calcium levels in DiSs. Our simulations indicate that a single inhibitory postsynaptic potential (IPSP) evoked in a DiS within 10 ms after an excitatory postsynaptic potential (EPSP) can curtail the local increase of calcium concentration without affecting the amplitude of membrane depolarization. This decoupling effect could impact long-term synaptic plasticity in cortical circuits.

## RESULTS

### Combining light and electron microscopy to access the ultrastructure of targeted populations of dendritic spines in brain slices

In the cortex, the morphology and distribution of dendritic spines vary depending on cortical area and layer in which the cell body is located [5,35,58,59], and dendritic spines are differently regulated depending on their location within dendritic trees — e.g. basal or apical dendrites [30,49,60–62]. Therefore, it is critical to take into account both the cellular and dendritic context to characterize the diversity of spine ultrastructure. To that aim, we developed a 3D-CLEM workflow allowing the ultrastructural characterization of dendritic spines on genetically-defined neuronal cell types and along identified types of dendrites in intact cortical circuits. In order to label specific subtypes of neurons, we used cortex-directed *in utero* electroporation (IUE) in mice. We electroporated neuronal progenitors generating layer 2/3 cortical PNs at embryonic day (E)15.5 with a plasmid expressing the fluorescent cytosolic filler tdTomato, granting access to the morphology of electroporated neurons, their dendrites and their dendritic spines in LM. We perfused adult mice with aldehyde fixatives, and collected vibratome sections of the electroporated area. To facilitate sample handling, we designed custom-made chambers allowing sample immersion in different solutions during confocal imaging and subsequent retrieval of the sample before EM preparation steps (S1 Fig). We enclosed 10-20 mm^2^ fragments of brain sections in these chambers and acquired images of optically isolated basal dendrites of bright electroporated neurons with confocal microscopy (Fig 1A).

**Fig 1.**
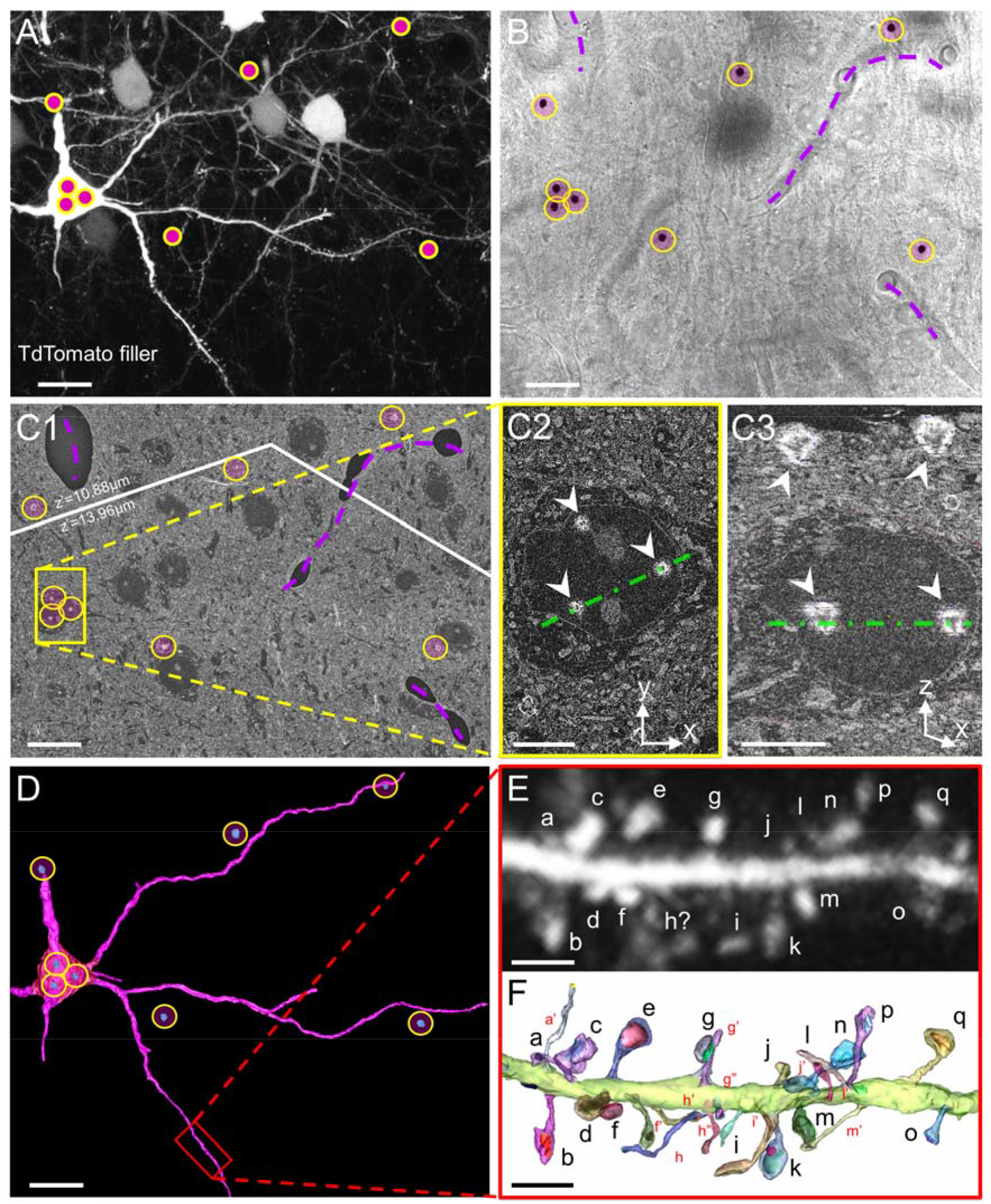
CLEM imaging of identified spines within intact cortical circuits. (A) Visualization of basal dendrites of a pyramidal neuron expressing cytosolic TdTomato in L2/3 of adult mouse SSC. DAB was photo-precipitated using focused UV light to insert correlative landmarks (pink dots in yellow circles). (B) Transmitted light image of the same field of view after DAB photo-precipitation. DAB precipitates are highlighted with yellow circles. Blood vessels are outlined with purple dashed lines. (C) Composite scanning EM (SEM) image displaying DAB patterning at the depth of the neuron of interest (yellow circles). Slight mismatch between LM and SEM observation planes resulted in DAB landmarks appearing in different z-planes during block-facing; the white line represents stitching between z-shifted images. In C1, landmarks are arranged as in B. C2 is a close-up on the soma of the electroporated neuron, labelled with three DAB landmarks (arrowheads). C3 corresponds to an orthogonal (x,z) view of the SEM stack along the green dashed line in C2. The superficial DAB layer enabled ROI targeting, and the deeper layer enabled retrospective identification of the target neuron. (D) 3D-reconstruction of dendrites of interest from the overview SEM stack. DAB landmarks are reconstructed in blue (in yellow circles). The red rectangle outlines the portion of dendrite represented in E and F. (E) Z-projection of the confocal stack corresponding to the portion of dendrite reconstructed in D. Letters identify individual spines. (F) 3D-EM reconstruction. Individual dendritic spines are manually segmented and randomly colored. Spines that were detected in EM but not in LM are labelled in red. Scale bars: A, B, C1, D: 10 μm; C2, C3: 5 μm; E, F: 2 μm.

A major challenge of CLEM in brain tissue is to recover the ROI in EM after imaging in LM. Several methods have been proposed to facilitate ROI recovery [50–52], but they come with caveats: (1) using only intrinsic landmarks has a low throughput [57,63]; (2) filling target neurons with 3,3-diaminobenzidine (DAB) masks intracellular ultrastructure [64]; (3) scarring the tissue with an infra-red laser to generate extrinsic landmarks, a.k.a. “NIRB” for “near-infrared branding” [56,65–68], produces landmarks with low pixel intensity in EM and can damage ultrastructure [63,69]. To facilitate ultrastructural measurements in non-obscured identified dendrites, we took advantage of the photo-oxidability of DAB [70,71]. We immersed the samples in DAB solution and applied focalized UV light at user-defined positions (Fig 1A) to imprint osmiophilic DAB landmarks around targeted dendrites (see panels B-E in S1 Fig) and pattern the tissue with localized electron-dense DAB precipitates (Fig 1B). After sample retrieval (see panel F in S1 Fig), tissue sections were processed for SBEM and embedded in minimal amounts of epoxy resin in order to maximize sample conductivity and SBEM image quality (see Methods). In 3D-EM stacks, ROIs were recovered within the complex environment of brain tissues using both intrinsic landmarks such as blood vessels (Fig 1B) and high-contrast DAB precipitates (Fig 1C; see also panel G in S1 Fig). We then segmented and reconstructed targeted dendrites in 3D, and registered whole portions of dendrites in both LM and EM to identify each dendritic spine unequivocally using neighboring spines as dependable topographic landmarks (Fig 1D). CLEM-based 3D-reconstruction enabled the identification of dendritic spines that were not visible in LM or EM alone. In LM, the limited axial resolution prevents the identification of axially oriented spines, which are easily detected in 3D-EM [49] (Fig 1D). On the other hand, spines with the longest and thinnest necks are conspicuous in LM stacks, but can be difficult to find in 3D-EM datasets without the cues provided by LM. The proportion of spines recovered with CLEM versus LM alone could amount to up to 30% per ROI, and 5% per ROI versus EM alone, highlighting the advantage of CLEM over unimodal microscopy approaches.

### Spine ultrastructure along the basal dendrites of L2/3 cortical pyramidal neurons

We used our CLEM workflow to quantify the full extent of the ultrastructural diversity of dendritic spines along the basal dendrites of layer 2/3 PNs of the SSC of three adult mice. We exhaustively segmented 254 μm of the basal dendritic arborization of four neurons and we reconstructed a total of 390 individual spines (S1 Table). As spine distance to the soma spanned from 20 to 140 μm, with basal dendrites extending up to 150 μm [72–75], our dataset can be considered representative of the whole spine population on these dendrites. The average linear density of dendritic spines was 1.5 ± 0.3 spine.μm^−1^. We then quantified the following parameters for each spine: neck length, neck diameter, head volume, head longitudinal diameter (referred to as “head length”), head orthogonal diameter (referred to as “head diameter”), number of PSDs, and PSD area (Fig 2A; S2 Table). In agreement with previous reports in both basal and apical dendrites of mouse cortical and hippocampal neurons [4,6,76,77], we found that ePSD area correlates linearly with the volume of the spine head (Fig 2B). We also observed a non-linear correlation between the length of the spine neck and its diameter (Fig 2C): long spines (neck length > 2 μm) always had thin necks (neck diameter < 0.2 μm), although short necks could also be thin. Furthermore, in spines with long necks, spine heads were always stretched longitudinally with respect to the neck (i.e. prolate) whereas they could also be stretched orthogonally (i.e. oblate) in shorter spines (Fig 2D), with a possible impact on nanoscale ion flows [78]. By contrast, there was no correlation between the position of the spine or the inter-spine distance and any of the morphological parameters we measured (S2 Fig). There was also no correlation between the length or the diameter of the neck and the morphometry of the spine head or ePSD (S1 Data), which is consistent with previous EM studies of L2/3 PNs of mouse neocortex [4,72] (but see [40,79] for different conclusions in other brain areas).

**Fig 2.**
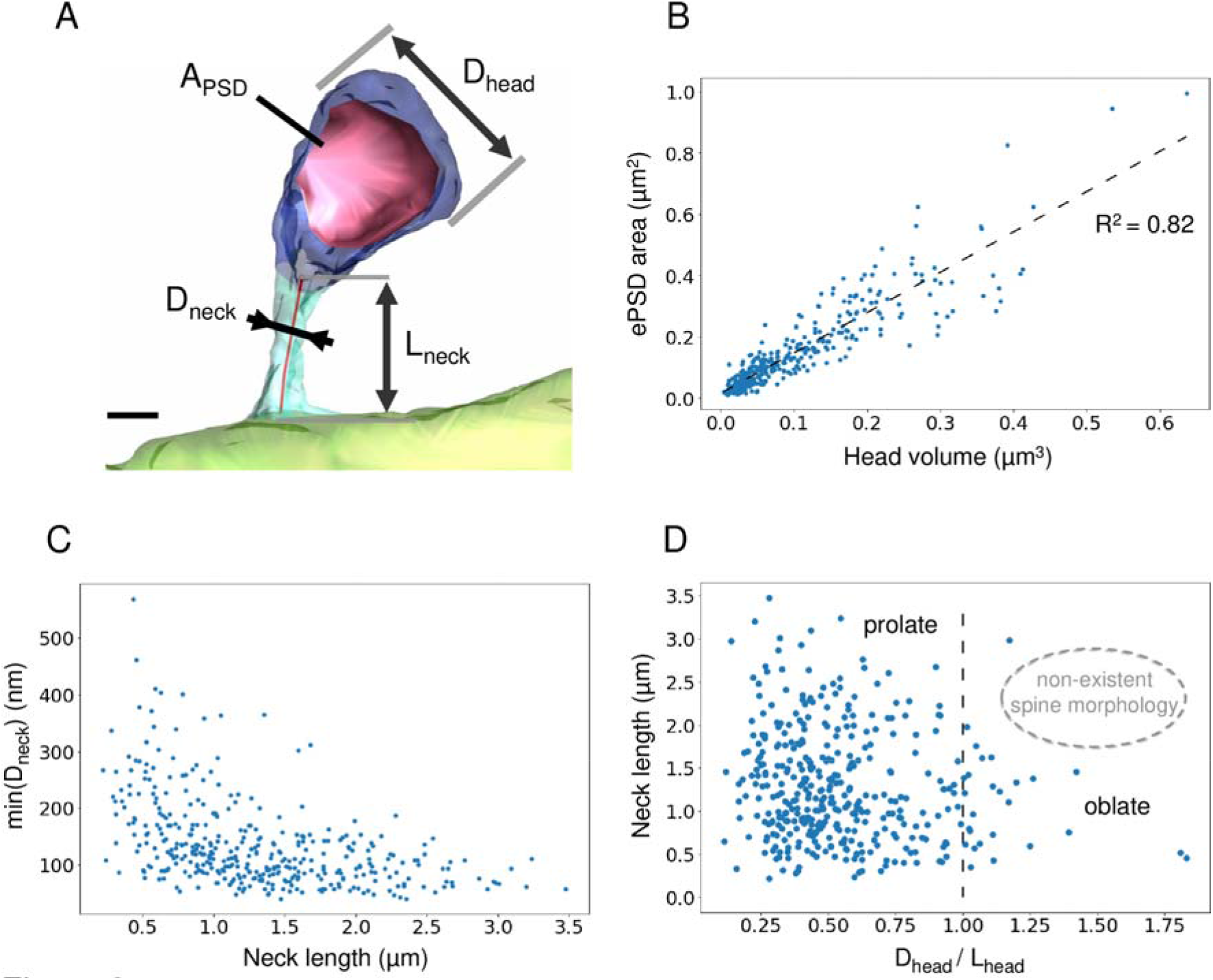
Spine morphometry along basal dendrites of layer 2/3 cortical pyramidal neurons. (A) 3D-reconstruction of a dendritic spine from a SBEM stack. Dendritic shaft is in light green, spine neck in turquoise, spine head in blue and PSD surface in red. The following parameters were measured : PSD area, head diameter, neck diameter and neck length. Scale bar: 300 nm. (B) Linear correlation of PSD area and spine head volume. R^2^=0.82. (C) Plot of the minimal spine neck diameter as a function of spine neck length. Spearman correlation coefficient is −0.58. (D) Neck length as a function of spine head orientation, as quantified by the ratio of spine head diameter (D_head_) over its length (L_head_). D_head_/L_head_ < 1 corresponds to a prolate spine head, which shape is stretched longitudinally with respect to the neck. D_head_/L_head_ > 1 characterizes an oblate spine head, oriented orthogonally to the neck. The dashed line corresponds to D_head_/L_head_ = 1. We did not observe any oblate spine with a long neck (non-existent spine morphology).

Since our CLEM approach grants access to the cytosolic content of spines (Fig 3A), we quantified the occurrence of SA, a complex stacked-membrane specialization of smooth endoplasmic reticulum (SER) which contributes to calcium signaling, integral membrane protein trafficking, local protein synthesis, and synaptic plasticity [12–14,80,81]. In basal dendrites, about 54% of spines contained a SA (Fig 3B), which is substantially higher than previous reports in the mature hippocampus [12,82]. These spines were randomly distributed along the dendrites. They had larger heads (Fig 3C), larger ePSDs (Fig 3D) and wider necks than spines devoid of SA (Fig 3E), consistent with previous morphological studies of CA1 PNs [12,82,83]. The probability that a spine contained a SA depending on spine head volume followed a sigmoid model (Fig 3F), predicting that all spines with a head diameter larger than 1.1 μm (21% spines in our reconstructions) contain a SA.

**Fig 3.**
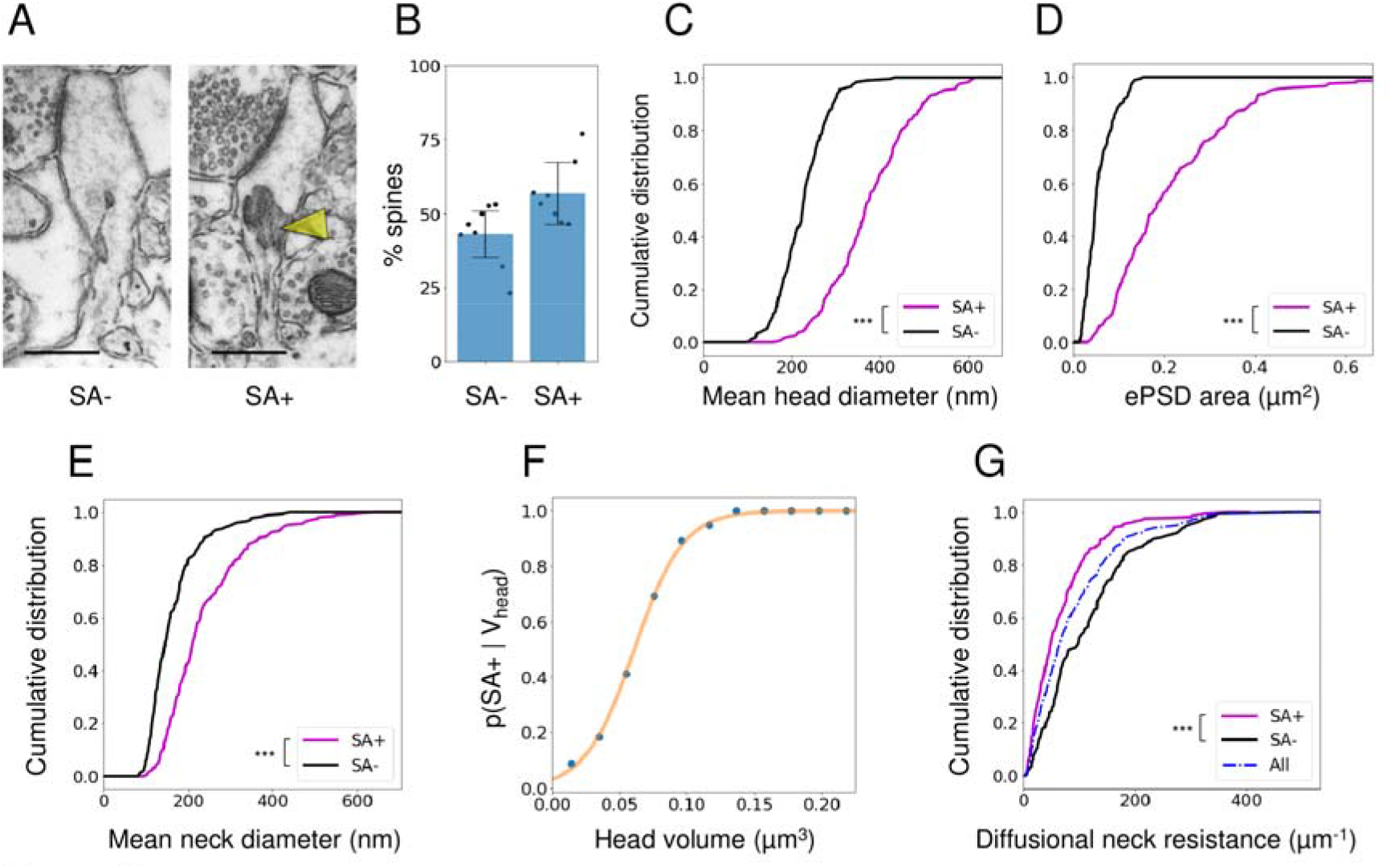
Ultrastructural comparison of spines with and without a spine apparatus. (A) TEM images of spines either devoid of spine apparatus (SA-, left) or containing a spine apparatus (SA+, right, yellow arrowhead). Scale bars: 500nm. (B) Proportion of SA- and SA+ spines. Histogram represents mean ± SD, from 390 spines in N=8 dendrites. (C) Distribution of mean head diameter for SA- and SA+ spines. N=179 and 221, respectively (p < 10^−38^). (D) Distribution of ePSD area. (p < 10^−40^) (E) Distribution of mean neck diameter. (p < 10^−12^) (F) Probability of harboring a SA as a function of spine head volume. Blue: experimental data. Orange: sigmoid fit. (G) Distribution of the diffusional resistance of the spine neck (W_neck_) calculated based on neck morphology. (p < 10^−5^). ***p < 0.001 calculated using Mann-Whitney test.

Next, we used our ultrastructural data to estimate the electrical resistance of spine necks using R_neck_ = ρW_neck_, where ρ is the cytosolic resistivity (set to 300 Ω.cm [84,85]) and W_neck_ is the diffusional neck resistance that restricts the diffusion of molecules and charges between spine heads and dendritic shafts [23]. To quantify W_neck_, for each spine we measured a series of orthogonal cross-sections of the neck along its principal axis and integrated W_neck_ = ∫ dℓ/ A(ℓ), where A(ℓ) is the neck cross-section area at the abscissa ℓ along the neck axis. W_neck_ ranged from 2 μm^−1^ to 480 μm^−1^ and R_neck_ from 8 MΩ to 1450 MΩ, with a median value of 188MΩ. These values are consistent with previous estimations based on EM reconstructions and STED super-resolutive light microscopy [17,86], and with direct electrophysiological recordings [87]. It has been proposed that the spine apparatus, which may occupy some of the spine neck volume, could increase W_neck_ [13,80,88]. Therefore, we subtracted SA cross-section from A(ℓ) when computing W_neck_ in SA+ spines (see Methods). This correction increased W_neck_ by 13% ± 2% in SA+ spines (S3 Fig). However, because of their wider necks, W_neck_ of SA+ spines was still lower (59% in average) than W_neck_ of spines devoid of SA (Fig 3G). These results suggest that, in addition to supplying large dendritic spines with essential resources, the SA may adjust W_neck_ and influence spine compartmentalization [12,13,82].

### Excitatory and inhibitory synapses in dually-innervated spines

We noticed that a small proportion of dendritic spines were contacted by two distinct pre-synaptic boutons (DiSs). DiSs have long been described in the literature as receiving both an excitatory and an inhibitory synaptic contact [89–92]. In the somato-sensory cortex, DiSs are contacted by VGLUT2-positive thalamocortical inputs [15] and they are sensitive to sensory experiences. The number of DiSs increases in response to sensory stimulation and decreases in response to sensory deprivation [73,93–95], suggesting their importance in synaptic integration and sensory processing. However, their scarcity in the cortex has been an obstacle to their ultrastructural and functional characterization. We took advantage of our CLEM approach and the molecular signature of this population of spines (i.e. the presence of a cluster of gephyrin, the core protein of inhibitory postsynaptic scaffolds [96,97]) to examine their morphological properties. To label inhibitory synapses in cortical PNs (Fig 4A), we co-expressed tdTomato with small amounts of GFP-tagged gephyrin (GFP-GPHN) [73,95,98,99]. We identified in LM spines containing a gephyrin cluster (Fig 4B) and we ascertained their dual-innervation in EM after back-correlating spine identity between LM and SBEM acquisitions. To do so, we aligned reconstructed dendrites on LM images (Fig 4C) and matched individual spines in both modalities (lettered in Fig 4B and Fig 4C). While ePSDs look asymmetrical and more electron-dense than inhibitory PSDs (iPSDs) in transmission EM [100,101], the anisotropic resolution of SBEM does not allow the distinction of ePSDs and iPSDs in most DiSs [49]. Therefore, we identified iPSDs on DiSs based on GFP-GPHN cluster position in LM images. In 89% of DiSs (33/37), the excitatory (GFP-GPHN-negative) PSD and the inhibitory (GFP-GPHN-positive) PSD could be clearly discriminated. However, in 11% of DiSs (4/37 DiSs), distinguishing ePSD from iPSD was not obvious due to the coarse axial resolution of LM imaging. To resolve ambiguities, we reconstructed the axons innervating the DiSs and determined their identity based on their other targets in the neuropil, either soma and dendritic shaft for inhibitory axons [2,102,103], or other dendritic spines for excitatory axons [49] (Fig 4D). As a result, we could unequivocally determine the excitatory or inhibitory nature of each synaptic contact on electroporated neurons, within ~10^5^ μm^3^ 3D-EM acquisition volume.

**Fig 4.**
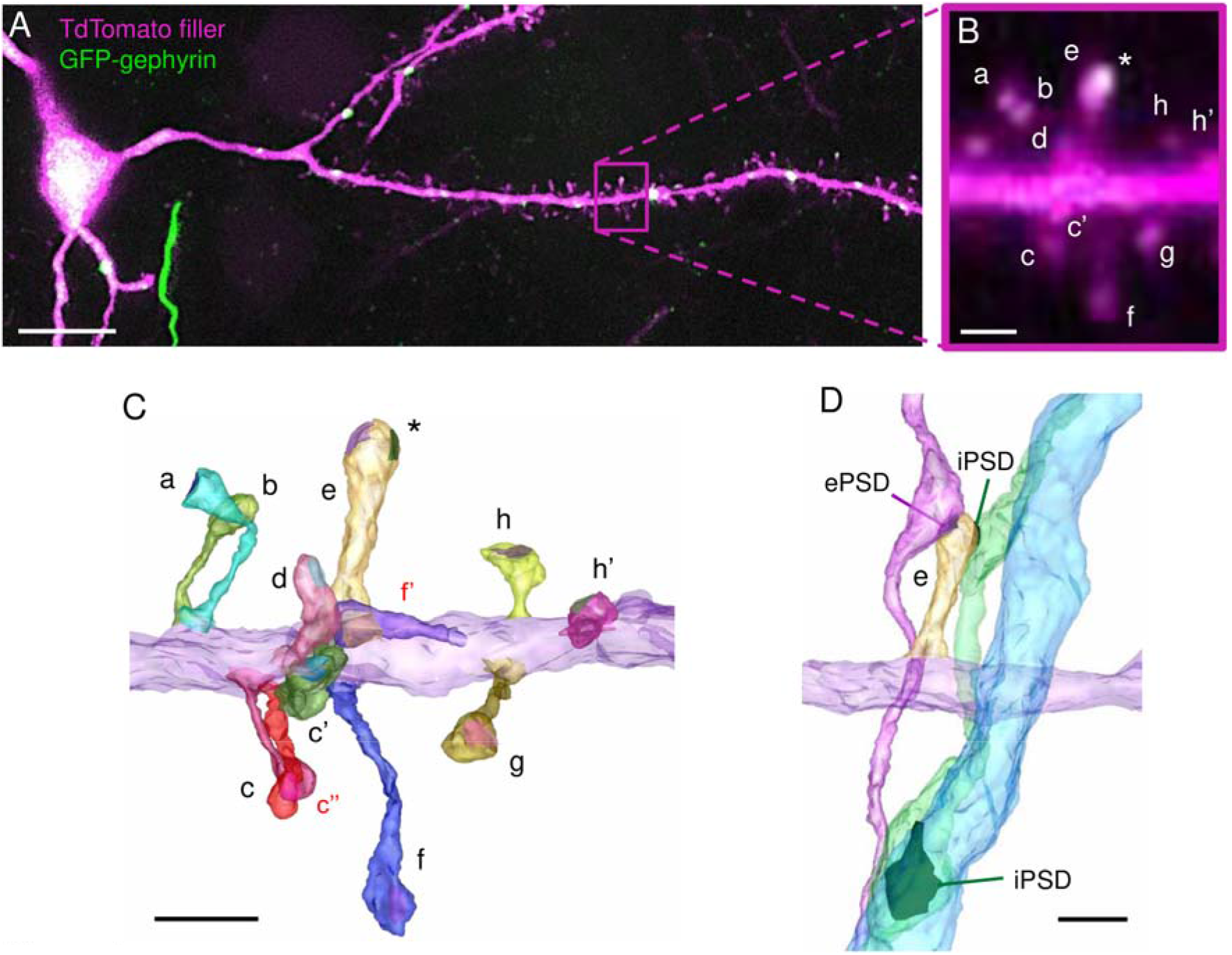
Identification of excitatory and inhibitory synapse on DiSs using CLEM. (A) Confocal image of basal dendrites of a cortical L2/3 PN that was electroporated with cytosolic TdTomato and GFP-GPHN to label inhibitory synapses. The magenta rectangle outlines the region enlarged in B. (B) Enlargement of a portion of the dendrite in A harboring several dendritic spine (lettered). Spine “e” contains a cluster of GFP-GPHN (asterisk) and corresponds to a putative dually-innervated spine (DiS). (C) 3D-EM reconstruction of the same dendritic fragment as in B. Dendritic shaft is colored in purple; individual spines and PSDs are colored randomly. Spines visible in EM but not in LM are labelled in red. The inhibitory PSD (colored in green) on spine “e” is identified based on the position of the GFP-GPHN cluster (asterisk in B and C). GFP-GPHN-negative PSDs are defined as excitatory. (D) 3D-EM reconstruction of spine “e” (yellow) with its presynaptic partners (magenta and green). As the “green” axon also targets a neighboring dendritic shaft (blue), it is defined as inhibitory. Scale bars: A: 10 μm; B, C, D: 1 μm.

In CLEM, we measured an average density of 1.4 ± 0.5 iPSDs per 10 μm of dendrite on DiSs and 2.1 ± 1.2 iPSDs per 10 μm of dendrite on the dendritic shaft— amounting to 3.5 ± 1.1 iPSDs per 10 μm of dendrite. iPSDs were homogeneously distributed either on spines or shaft from 24 μm away from the soma to the dendritic tip, which contrasts with apical dendrites where spinous inhibitory synapses are distally enriched [73]. Along the basal dendrites of L2/3 cortical PNs, 38% of inhibitory contacts occurred on dendritic spines, which is higher than previously estimated using LM only [73,99]. DiSs represented 10% ± 3% of all spines (Fig 5A). They had larger heads than SiSs (Fig 5B), in line with previous reports [15,104], and 86% ± 13% of them contained a SA (Fig 5C). DiSs also differed in terms of neck morphology. They had longer necks than SiSs of comparable head volume (V_head_ > 0.05 μm^3^), although neck length distribution was similar in the whole populations of SiSs and DiSs (Fig 5D). DiSs also had lower D_neck_/V_head_ ratio than SiSs (Fig 5E), although D_neck_ distribution was similar between SiSs and DiSs (S4 Fig), suggesting that excitatory signals generated in DiSs are more compartmentalized than signals of similar amplitude generated in SiSs. Accordingly, DiSs had a higher W_neck_ than SiSs of comparable head size (52% larger in average) (Fig 5F). In spine heads, ePSDs on DiSs were larger than ePSDs on SiSs (174% ± 113% of ePSD area) (Fig 5G), consistent with the larger head size of DiSs. By contrast, iPSDs on DiSs were smaller than shaft iPSDs (53% ± 15% of shaft iPSD area) (Fig 5H). The area of iPSDs on DiSs did not correlate with spine head volume (S5 Fig). In 95% of DiSs, iPSDs were smaller than ePSDs (half the area, in average) (Fig 5I). Together, these results indicate that DiSs represent a specific population of dendritic spines with distinctive ultrastructural features that could impact their functional properties.

**Fig 5.**
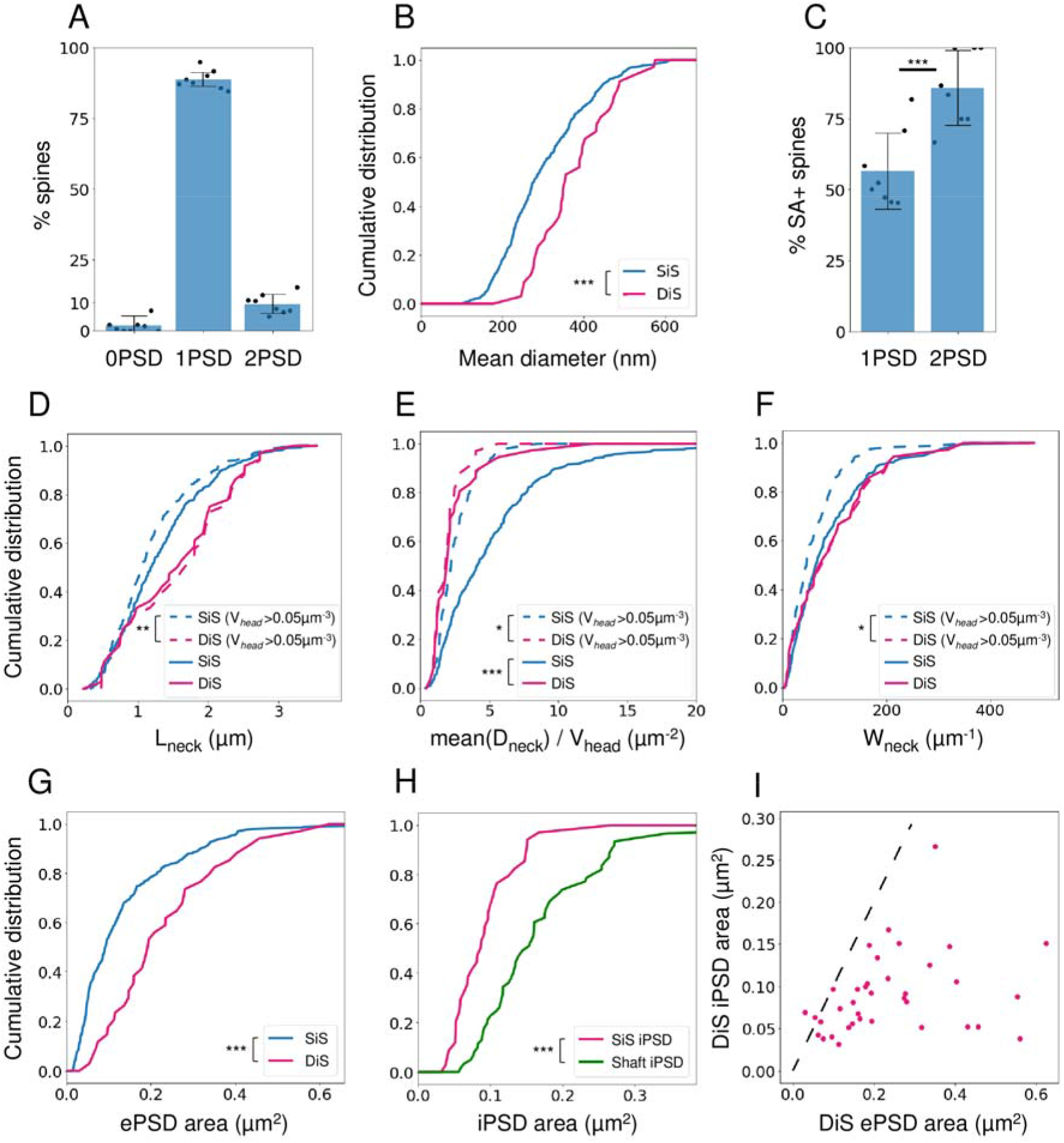
Anatomical properties of DiSs. (A) Proportion of spines harboring 0, 1 or 2 synaptic contacts, quantified with CLEM. Histograms represent mean ± SD, from 390 spines in N=8 dendrites. (B) Quantification of mean spine head diameter for SiSs (blue) and DiSs (red). (p < 10^−4^). (C) Proportion of SiSs and DiSs harboring a SA. (p < 10^−10^ using Pearson’s χ^2^ test) (D-F) Quantification of neck length (D), the ratio between mean neck diameter and head volume (E) and the diffusional neck resistance (W_neck_) (F) between SiSs and DiSs (solid lines, N=349 and 37, respectively) and between DiSs with SiSs of similar head volume (spines with V_head_ > 0.05 μm^3^, dashed lines, N=186 and 34, respectively). (G) Quantification of iPSD area in DiSs and dendritic shafts. N=37 and 62, respectively (p < 10^−6^). (H) Quantification of ePSD area in SiSs or DiSs. (p < 10^−5^). (I) Plot of iPSD area as a function of ePSD area in individual DiSs. The dashed line (y = x) highlights that the ePSD is larger than the iPSD in most of DiSs. N=37. p-values were computed using Mann-Whitney test (B, D-H) or Pearson’s χ^2^ test (C). Only significant (p < 0.05) p-values are shown (*p < 0.05; **p < 0.01; ***p < 0.001).

### Morphologically constrained model of synaptic signaling

Next, we wanted to assess the impact of spine diversity on synaptic signals. We used a computational approach based on a multi-compartment “ball-and-stick” model of the neuronal membrane [40,105]. This model comprises an isopotential soma and two dendritic compartments structured as cables featuring passive resistor-capacitor (RC) circuits and conductance-based synapses. The two dendritic compartments correspond to the dendrite receiving the synaptic inputs and to the remainder of the dendritic tree (Fig 6A1) [106,107]. We constrained this model with morphological parameters measured in CLEM (i.e. distance between spine and soma, neck resistance, head volume, head membrane area, ePSD area and iPSD area for 390 spines, and dendritic diameter), taking into account the structural shrinkage resulting from chemical fixation (S6 Fig). Excitatory and inhibitory synaptic conductances were modeled as bi-exponential functions of time, with their rise and decay times tuned to the kinetics of different receptor types: AMPA and NMDA receptors at ePSDs, and type A γ-aminobutyric acid (GABA_A_) receptors at iPSDs (Fig 6A2; see Methods). Individual synaptic conductances were scaled proportionally to PSD areas [9,77,108]. Voltage-dependent calcium channels (VDCCs) in spine heads were modeled using Goldman–Hodgkin–Katz equations [109] and their conductance was scaled proportionally to spine head areas. We adjusted excitatory synaptic conductivity so that average amplitudes of both synaptic currents and somatic depolarizations evoked by individual excitatory postsynaptic potentials (EPSPs) fitted published electrophysiological values [110–113] (see Methods). After calibration of excitatory synapses, maximal synaptic conductance (g) ranged from 0.04 nS to 3.13 nS for g_AMPA_ (0.456 ± 0.434 nS) and from 0.04 nS to 3.42 nS for g_NMDA_ (0.498 ± 0.474 nS), in line with the literature [114]. We then adjusted inhibitory synaptic conductivity to set the mean conductance of dendritic inhibitory synapses to 1 nS [37,115,116]. As a result, g_GABA_ ranged from 0.33 nS to 3.36 nS (1.00 ± 0.577 nS) for synapses located on the shaft, and from 0.19 nS to 1.56 nS (0.528 ± 0.277 nS) for inhibitory synapses located on spines.

**Fig 6.**
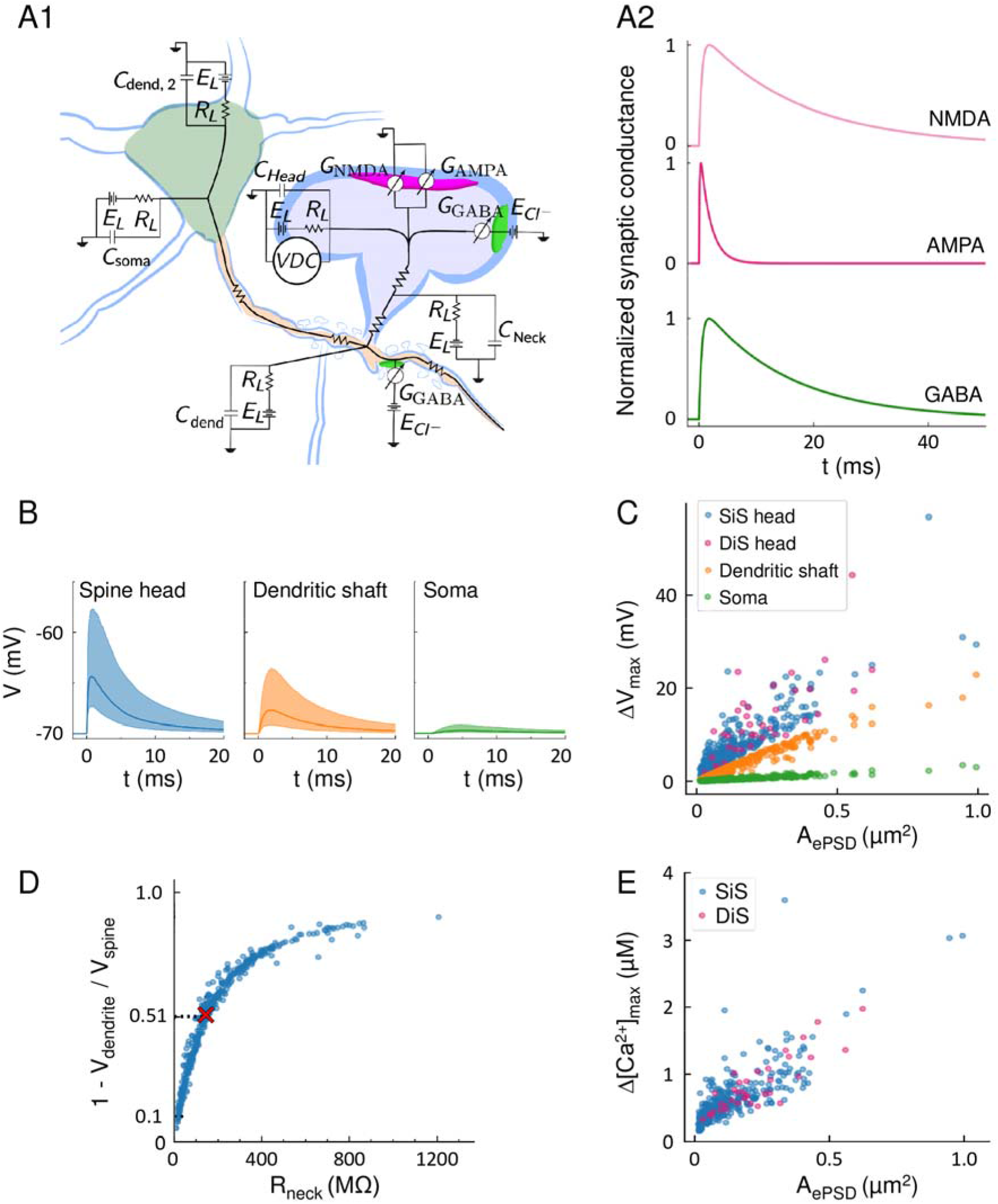
Morphologically-constrained model of synaptic signaling. (A) Schematic of the circuit model (A1) and representative time-course of excitatory (magenta) and inhibitory (green) conductance based on the kinetics of AMPA, NMDA and GABA_A_ receptors (A2). All compartments include passive resistor-capacitor circuits to model cell membrane properties and optionally include an active conductance that models voltage-dependent currents (VDC). All modeled spines feature an excitatory synapse with glutamatergic AMPA and NMDA currents. Spines and dendritic compartments can also feature an inhibitory synapse with GABAergic currents. All conductances were scaled to PSD area (see Methods). (B) Simulation of the time-courses of membrane depolarization following an EPSP, taking into account spine diversity (i.e. R_neck_, ePSD area and distance to soma, as measured in CLEM). Membrane voltage is monitored over time in the spine head (blue), in the dendritic shaft in front of the spine (orange) and in the soma (green). Median voltage transients are plotted as solid lines. Shaded areas represent 68% confidence intervals, which span approximately one standard deviation on each side of the mean. (C) Amplitude of evoked depolarization (ΔV_max_) as a function of ePSD area at three distinct locations: head of SiSs (blue) or DiSs (magenta) where the EPSP was elicited, dendritic shaft 1 μm from the spine (orange) or soma (green). (D) Attenuation of the amplitude of depolarization between the spine head and the dendrite as a function of the resistance of the neck (R_neck_). The attenuation was calculated as: α = 1 − ΔV_max_, shaft /ΔV_max,spine_. Red cross: mean value of α. (E) Estimated amplitude of intracellular calcium concentration transients Δ[Ca^2+^]_max_, following activation of NMDA receptors and VDCCs as a function of ePSD area. Three spiking outliers are not represented.

We first examined the propagation of simulated EPSPs. We compared the evoked depolarization amplitude ΔV_max_ in three different compartments: in the spine head in which the EPSP was elicited, in the dendritic shaft close to the spine, and in the soma (Fig 6B). ΔV_max_ followed a log-normal distribution, reflecting the morphological variability of spines (Fig 6B). Due to the passive attenuation of electrical signals along dendritic processes, ΔV_max_ was sharply attenuated between the head of the spine and the dendritic shaft (51% attenuation in average) and about 5% of ΔV_max_ reached the soma (Fig 6B), in line with measurements performed in basal dendrites of L5 cortical PNs using voltage dyes, electrophysiology and glutamate uncaging [25,117]. ΔV_max_ scaled with ePSD area in all compartments (Fig 6C). To determine the contribution of morphological parameters to the variance of ΔV_max_, we used a generalized linear model (GLM) [118]. We analyzed the contribution of the volume, diameter and resistance of spine necks and heads as well as the contribution of ePSD area and distance between spine and soma (L_dend_) to the amplitude of the signals in the soma and in spine heads. In the soma, ΔV_max_ was mainly determined by A_ePSD_, which accounted for 89% of its variance when EPSPs were elicited in SiSs (77% when they came from DiSs). The second determinant was L_dend_, which accounted for 6.4% of the variance of ΔV_max_ for EPSPs generated in SiSs (14.7% in DiSs). The contribution of R_neck_ to the variance of ΔV_max_ in the soma was comparatively negligible (S3 Table), indicating that the passive attenuation of EPSPs along dendrites dominates the contribution of R_neck_ to somatic depolarizations evoked in spines. In the heads of SiSs, A_ePSD_ and R_neck_ accounted for 60% and 19% of the variance of ΔV_max_, respectively (also see S7 Fig for the dependence of ΔV_max_ on R_neck_). In the heads of DiSs, the contribution of R_neck_ to ΔV_max_ was much higher, reaching 38% of the variance, while A_ePSD_ contribution dropped to 47% (S3 Table). In 56% of dendritic spines, R_neck_ was large enough (>145MΩ) to attenuate EPSP amplitude by >50% across the spine neck, and more than 90% of spine necks attenuated the signal by at least 10% (Fig 6D), suggesting that most spine necks constitutively compartmentalize electrical signals in the head of spines.

We also estimated the elevation of calcium ion concentration (Δ[Ca^2+^]) in spine heads induced by an EPSP. Δ[Ca^2+^] was similar in SiSs and DiSs and varied non-linearly with A_ePSD_ (Fig 6E). A_ePSD_ was the main determinant of Δ[Ca^2+^], accounting for 30% of the variance in SiSs (45% in DiSs), followed by R_neck_ (9%; S3 Table). As a single EPSP is not sufficient to elicit a Ca^2+^ spike, we did not model Ca^2+^ transients outside of spine heads. Overall, our model provides quantitative insights into the variability of EPSP amplitude originating from spine diversity and highlights differences in the contribution of morphological parameters to spine depolarization and calcium signals in DiSs and SiSs.

### Spatial interplay of excitatory and inhibitory signals

We used our model to compare the effects of spinous and dendritic shaft inhibition. *In vitro* uncaging experiments have shown that inhibitory contacts located on DiSs could weaken local calcium signals [119], but the consequences on synaptic excitation are still unclear [75,120]. To understand how spine ultrastructure and iPSD location influence synaptic integration, we modeled the interaction between one IPSP and one EPSP under the constraint of our morphological measurements. Assessing the extent of signal variability originating from spine morphological heterogeneity requires a large number of simulations (N≥1000). Therefore, we used a bootstrapping method [121] (see Methods) to derive N≥1000 sets of parameters from our dataset of 390 spines and 62 shaft iPSDs (S1 Data), and provide unbiased estimations of the mean and variance of the signals. Importantly, the strength of inhibition depends on the reversal potential of chloride ions (E_Cl_^−^). In healthy mature layer 2/3 cortical PNs, E_Cl_^−^ typically varies between the resting membrane potential, V_rest_ = −70 mV, and hyperpolarized values (−80 mV) [122]. When E_Cl_^−^ = −70 mV, active inhibitory synapses generate a local increase of membrane conductance, which “shunts” membrane depolarization induced by concomitant EPSPs. When E_Cl_^−^ < −70 mV, the driving force of Cl^−^ ions is stronger, GABAergic inputs can hyperpolarize the cell membrane and IPSPs counter EPSPs. These two situations are respectively termed “shunting inhibition” and “hyperpolarizing inhibition” [37,122].

We first assessed the impact of “shunting” IPSPs elicited in the dendritic shaft on individual EPSPs depending on the inter-synaptic distance (Fig 7A). For each iteration of the model (i.e. N=3700 sets of realistic morphological values for spine size, and iPSD location and size), we varied the distance Δx between a spine receiving an EPSP and a shaft iPSD activated simultaneously. For Δx > 0, the iPSD was located between the spine and the soma (i.e. “on-path” inhibition, from the viewpoint of the soma), and for Δx < 0, the iPSD was located distally to the spine (i.e. “off-path” inhibition). To quantify inhibition, we compared the amplitude of individual EPSPs in the absence (ΔV_max,E_) or presence (ΔV_max,E+I_) of inhibition, and computed the drop in depolarization amplitude *inh_V_* = 1 − Δ_max,E+I_ / ΔV_max,E_ . *inh_V_* = 0 indicates that the electrical signal was not affected, *inh_V_* = 1 indicates that it was completely inhibited. Importantly, due to the electrical properties of the cell membrane, *inh_V_* depends on where the signal is measured. In the soma, *inh_V_* was maximal (20% in average) when the iPSD was located on-path and it decreased exponentially with Δx when the iPSD was located off-path (exponential decay length: L_soma,off_ = 30 μm) (Fig 7B), highlighting the impact of proximal inhibitory synapses on distal excitatory inputs [37,123]. In the spine where the EPSP was elicited, *inh_V_* decayed exponentially with Δx for both on-path and off-path inhibition (Fig 7C), with respective decay lengths L_spine,on_ = 30 μm and L_spine,off_ = 11 μm (also see S8 Fig for the dependence of decay lengths on the dendritic cross-section). Therefore, shaft inhibition could affect excitatory signals within 40 μm, which corresponds to approximately 50 spines considering the spine density and dendritic diameter that we measured in basal dendrites of L2/3 cortical PN.

**Fig 7.**
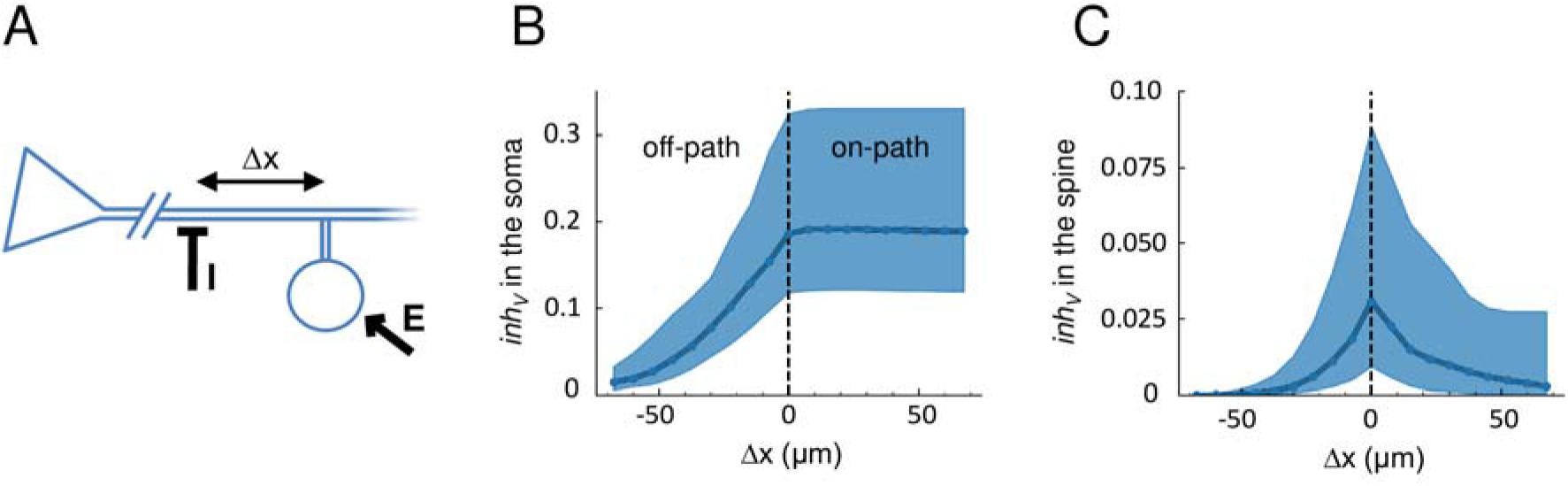
Effect of the distance between excitatory and inhibitory synapses on the integration of coincident EPSPs and IPSPs. (A) Sketch: an EPSP was elicited in the spine and an IPSP in the shaft at a distance Δx from the spine. (B-C) Voltage inhibition, *inh_V_*, calculated in the soma (B) or in the spine head (C) as a function of Δx, for N=3700 iterations of the model. On-path inhibition: Δx > 0; off-path inhibition: Δx < 0. Solid lines represent medians. Shaded areas represent 68% confidence intervals, which span approximately one standard deviation on each side of the mean.

Next, we focused on DiSs. We modeled N=3700 DiSs and shaft iPSDs (Δx = 0) at random locations on the dendrite, and we compared two configurations per iteration of the model: (1) the ePSD of the DiS and the shaft iPSD were activated simultaneously (Fig 8A1); (2) both the ePSD and the iPSD of the DiS were activated simultaneously (Fig 8A2). In the soma, the effect of shaft and spinous inhibition was comparable: *inh_V_* was centered on 17%, and reached up to 50% (Fig 8B), in line with somatic recordings following coincident uncaging of glutamate and GABA in acute brain slices [119]. By contrast, in the head of all DiSs, spinous inhibition was more efficient than shaft inhibition despite the smaller size of spinous iPSDs compared to shaft iPSDs. Spinous *inh_V_* was centered on 10% and reached up to 35%, whereas dendritic *inh_V_* was centered on 3% and did not reach more than 23% (Fig 8C). These results support the notion that the placement of inhibitory synapses structures the detection and integration of excitatory signals [103,124,125], and highlight the role of spinous inhibition in local synaptic signaling.

**Fig 8.**
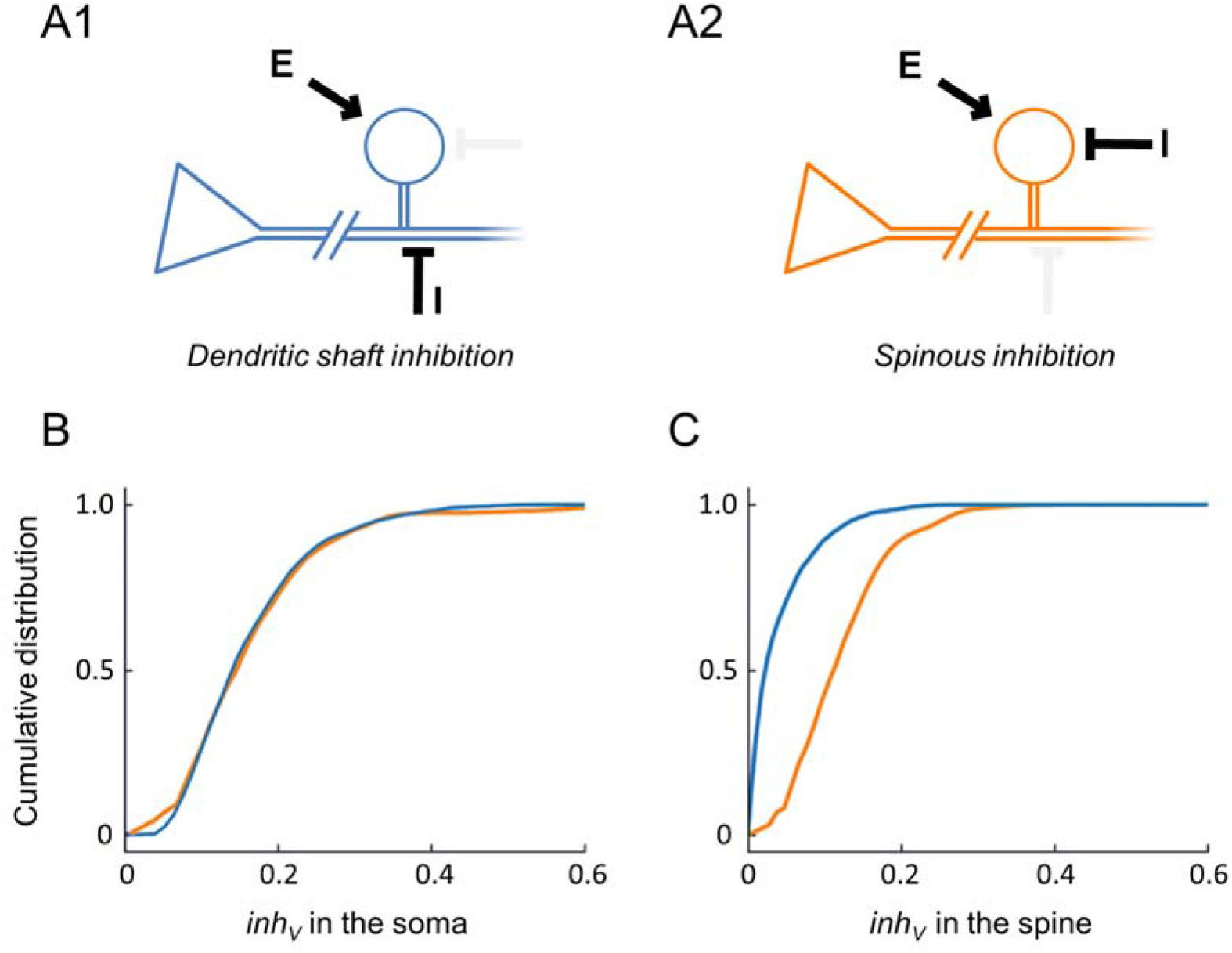
Effect of dendritic and spinous inhibition on EPSPs. (A) Sketches: an EPSP (arrow) was elicited in a bootstrapped DiS placed randomly along the dendrite, and an IPSP (┬ symbol) was elicited either in the dendritic shaft at Δx = 0.7 μm from the stem of the spine (A1, blue) or directly in the spine head (A2, orange). (B-C) Quantification of the inhibitory impact, *inh_v_*, in the soma (B) and in the spine head (C) for N=3700 iterations of the model. Blue: dendritic inhibition; Orange: spinous inhibition.

We then run the simulations with a lower chloride reversal potential (E_Cl_^−^ = −80 mV) for which GABAergic inputs can hyperpolarize the cell membrane. In numerous simulations, we observed that hyperpolarization could take over depolarization (i.e. *inh_V_* > 1) (see panel A in S9 Fig). This was the case in the soma for 37% of simulations, and in the spine where the EPSP was generated for 10% of simulations. With E_Cl_^−^ = −80 mV, the median *inh_V_* imposed by spinous inhibition in spine heads was 45%, in line with a previous morphologically constrained model [75] and with the fact that a hyperpolarizing IPSP may block multiple

EPSPs depending on iPSD placement [103]. Interestingly, lowering E_Cl_^−^ affected spinous and shaft inhibition differently. In the soma, shaft inhibition became more efficient than spinous inhibition (see panel B in S9 Fig) due to the larger area of shaft iPSDs. However, in the heads of most DiSs, spinous inhibition remained more efficient than shaft inhibition (see panels C-F in S9 Fig). Altogether, our results indicate that spinous inhibition is stronger than shaft inhibition in DiSs, and that their relative weight is modulated by the driving force of chloride ions.

### Temporal interplay of excitatory and inhibitory signals

Since the efficacy of inhibition depends on the membrane potential at the onset of the IPSP [36,38,123,126], we addressed the effect of input timing on inhibition efficacy in DiSs. We simulated the interaction of one EPSP and one IPSP generated with a time difference of Δt (Fig 9A). For Δt < 0 (IPSP before EPSP), IPSPs decreased ΔV_max_ (Fig 9B1). For Δt > 0 (IPSP after EPSP), IPSPs had no effect on ΔV_max_ but abruptly decreased the tail of the EPSPs [126] (Fig 9B2). We first compared how spinous and shaft inhibition reduced EPSP duration by comparing the 80-to-20% decay time of the summed signals (τ_E+I_) to that of uninhibited EPSPs (τ_E_). Decay times were minimal at Δt = +4 ms and decreased by 64% and 78% with spinous and shaft inhibition respectively (Fig 9C), which could shorten the integration window of EPSPs and increase the temporal precision of synaptic transmission [127]. The variance of τ_E_ was mainly determined by the area of iPSDs (S4 Table). We then quantified how the timing of inhibition affected EPSP amplitude using *inh_V_* (Δt) = 1 − ΔV_max_,_E+I_ / ΔV_max,E_. In spine heads, *inh_V_* was an asymmetrical function of Δt [36,126] and it was maximal at Δt = −4 ms for spinous inhibition, and at Δt = −6 ms for dendritic shaft inhibition. Overall, spinous inhibition was stronger than shaft inhibition, decreasing ΔV_max_ by 26.3% and 16.2%, respectively (median values in Fig 9D; see also panel A in S10 Fig for *inh_V_* (Δt) with higher E_Cl_^−^). Interestingly, R_neck_ had a negligible contribution to the variance of *inh_V_* in the case of hyperpolarizing inhibition, but a major one (37%) in the case of shunting inhibition (S4 Table), suggesting that spine necks compartmentalize IPSPs differently depending on E_Cl_^−^.

**Fig 9.**
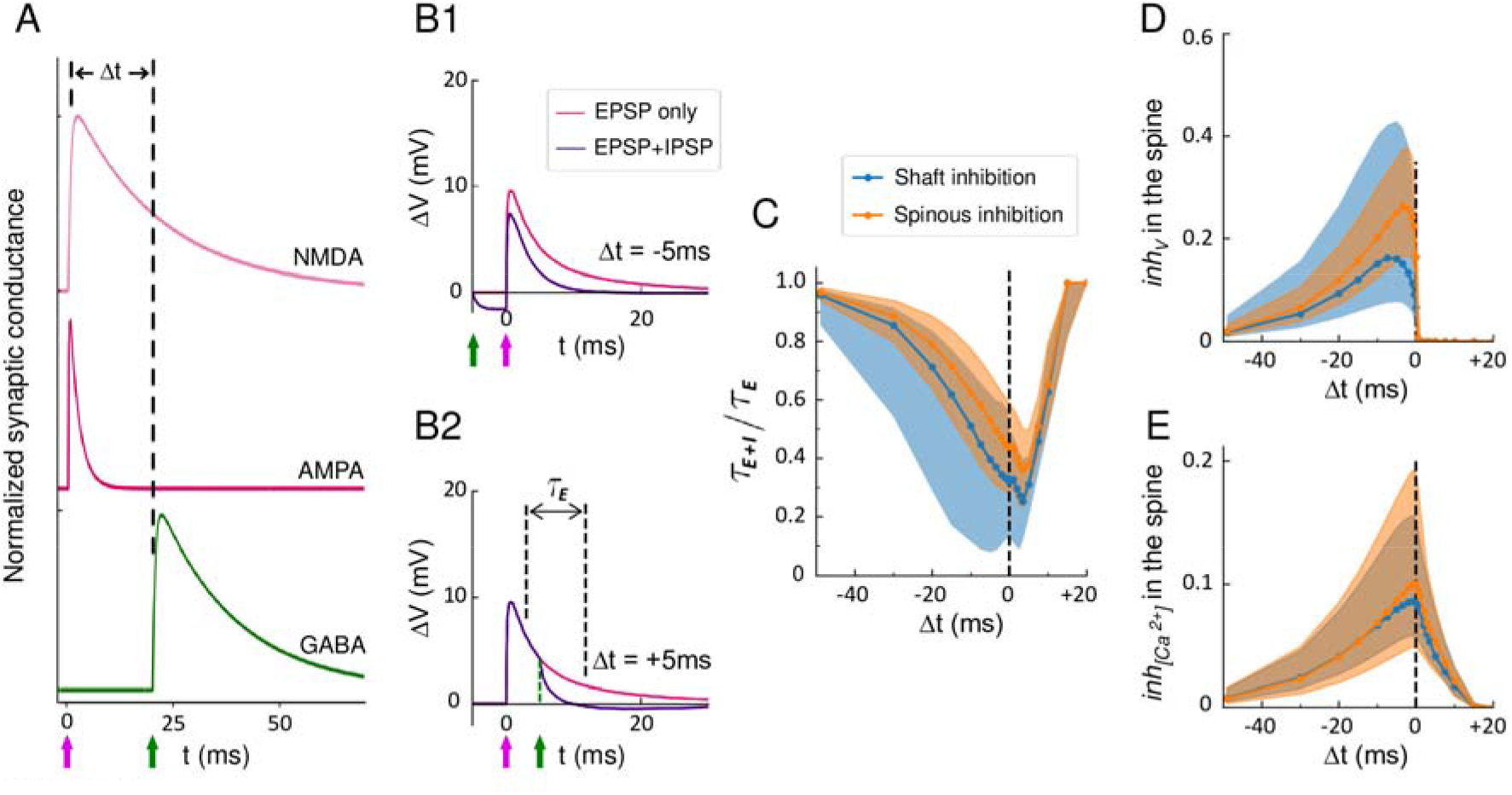
Effect of input timing on EPSP and IPSP integration. (A) Schematic: excitatory AMPA and NMDA conductances were activated at t=0. The inhibitory GABAergic conductance was activated at an interval Δt before or after the onset of excitation. (B) Examples of the time-course of depolarization in the spine head for Δt = +5 ms (B1) and Δt = −5 ms (B2) (purple curves) compared to no inhibition (magenta curves) for E_Cl_^−^ = −80 mV and V_rest_ = −70 mV. Arrows represent the onset of excitatory and inhibitory inputs (magenta and green arrows, respectively). τ_E_ represents the 80-to-20% decay time (case with no inhibition). (C) Ratio of 80-to-20% decay time of membrane depolarizations in the presence of inhibition (τ_E+I_) over that without inhibition (τ_E_) as a function of Δt for dendritic (blue) or spinous (orange) inhibition. Solid lines represent medians. Shaded areas represent 68% confidence intervals, which span approximately one standard deviation on each side of the mean. (D) Voltage inhibition in the spine head, *inh_V_*, induced by dendritic (blue) or spinous (orange) IPSPs as a function of Δt. (E) Inhibition of the calcium influx in the spine head, *inh_[Ca^2+^]_*, induced by dendritic (blue) or spinous (orange) IPSPs as a function of Δt.

Then, we examined the effect of timed inhibition on calcium signaling in spines, using *inh_[Ca^2+^]_* (Δt) = 1 – [Ca^2+^]_max_,_E+I_ / [Ca^2+^]_max,E_. We observed that *inh_[Ca^2+^]_* peaked at Δt = 0 ms for both spinous and shaft inhibition. More precisely, spinous inhibition reduced calcium transient amplitude by 10% in average, reaching >36% in the top 10% simulations, while shaft inhibition reduced it by 8.6% in average and >28% in the top 10% simulations (Fig 9E). These values are in the range of *inh_[Ca^2+^]_* measured with double uncaging experiments [119]. Importantly, IPSPs could decrease the amplitude of calcium transients, within a short time-window (Δt between 0 and +10 ms) in which depolarization amplitude was not affected (Fig 9D-E), thereby decoupling calcium signalling from electrical activity in DiSs.

## DISCUSSION

In the present study, we developed a novel 3D-CLEM workflow allowing the ultrastructural characterization of specific populations of dendritic spines in genetically defined types of neurons. We used this workflow to exhaustively reconstruct spines and synaptic contacts along the basal dendrites of fluorescently labelled L2/3 cortical PNs of the SSC and to provide a quantitative description of their diversity. We input our measurements in a computational model to analyze the variability of electrical and calcium synaptic signals originating from spine ultrastructural diversity, and to characterize the spatio-temporal integration of excitatory and inhibitory inputs. Our results shed light on unique properties of DiSs, which represent 10% of all spines and 38% of all inhibitory synapses along the basal dendrites of L2/3 cortical PNs. We show that while individual inhibitory synapses distributed along dendritic shafts can be powerful enough to block several EPSPs, spinous inhibitory synapses affect excitatory signals more efficiently in DiSs. We also show that the activation of a spinous inhibitory synapse within a few milliseconds after an EPSP can decouple voltage and calcium signals in DiSs, which could impact calcium-dependent signaling cascades that drive spine plasticity.

The molecular composition of synapses and their biophysical properties are reportedly heterogeneous along dendrites and across dendritic trees [35,39,128,129]. However, most computational models so far have used *ad hoc* or averaged values as parameters for dendritic spines and excitatory synapses [130–132], and considered that all inhibitory synapses were located along the dendritic shaft [133]. The correlative approach we propose provides an accessible solution for detailed quantification of synaptic diversity beyond the μm scale in intact brain circuits, which may help improve the accuracy of computational models. Our workflow is applicable to any type of tissue and allows anatomical measurements of any kind of genetically labelled cells and organelles. One technical limitation is the need for chemical fixation, which may distort tissue morphology [134,135]. Therefore, it may be necessary to correct for tissue shrinkage based on a morphological comparison with physically fixed tissues (see panel B3 in S6 Fig) in order to reliably depict *in vivo* situations. Future development of aldehyde-free cryo-CLEM methods will be important to grant access to cellular and synaptic ultrastructure in close-to-native environments.

Applying 3D-CLEM to the basal dendrites of L2/3 cortical PNs allowed us to quantitatively describe the landscape of synaptic diversity and to characterize the ultrastructural features of a scarce population of dendritic spines receiving both excitatory and inhibitory synaptic inputs (DiSs). In the cortex, DiSs are mostly contacted by VGluT2-positive excitatory thalamo-cortical inputs [15] and they receive inhibition from somatostatin-expressing and parvalbumin-expressing interneurons [119,136], which are the two main sources of inhibitory inputs to the basal dendrites of layer 2/3 cortical PNs [137,138]. *In vivo* 2-photon imaging experiments have shown that DiSs are among the most stable spines along the dendrites of layer 2/3 PNs [104]. The inhibitory synapse in DiSs is smaller and more labile than inhibitory synapses along dendritic shafts, and it is very sensitive to sensory experience [73,94,95,104]. Whisker stimulation induces a lasting increase in the occurrence of iPDSs in spines of the barrel cortex [94] and monocular deprivation destabilizes iPSDs housed in spines of the visual cortex [73,95,104], suggesting their role in experience-dependent plasticity. Our morphological and computational analysis provides new insights into the biophysical properties of DiSs. We show that DiSs have larger heads and larger ePSDs than SiSs, and most often contain a spine apparatus. However, the ratio between mean spine neck diameter and spine head volume (or ePSD area) was smaller in DiSs than in SiSs, and DiSs had longer necks than SiSs of comparable head volume, so that EPSPs of similar amplitudes encounter a higher neck resistance in DiSs than in SiSs. Thus, DiSs are uniquely compartmentalized by their ultrastructural features and the presence of an inhibitory synapse.

Our model predicts that IPSPs occurring in DiSs within milliseconds after an EPSP can curtail calcium transients without affecting depolarization, thereby locally decoupling voltage and calcium signaling. This is expected to impact the induction of long-term forms of synaptic plasticity, such as long-term potentiation (LTP) or long-term depression (LTD), which underlie learning and memory [80,139–141]. The induction of LTP versus LTD is determined by the magnitude and time course of calcium flux, with brief, high calcium elevation generating LTP, sustained moderate calcium elevation generating LTD, and low calcium level inducing no plasticity [142–144]. Therefore, a small reduction in the amplitude of calcium transients may limit spine potentiation, or even cause depression [120,145,146]. In the cortex, thalamocortical inputs may contact DiSs on the basal dendrites of L2/3 both directly (excitatory connection) and indirectly through feed-forward inhibition *via* parvalbumin-expressing fast-spiking interneurons [127,147]. The delay between thalamo-cortical excitatory and feed-forward inhibitory signals is typically +1 ms to +3 ms [127], within the 10 ms time window for voltage-calcium decoupling in DiSs. Therefore, the presence of inhibitory synapses in DiSs could prevent synaptic potentiation and thereby increase the temporal precision of cortical response to sensory stimulation [94,127,148]. On the contrary, the removal of spine inhibitory synapses could favor synaptic potentiation during experience-dependent plasticity such as monocular deprivation to strengthen inputs from the non-deprived eye [73,95,149].

Our understanding of synaptic and dendritic computations is intimately linked to the quantitative description of synaptic distribution, ultrastructure, nano-organization, activity and diversity in neural circuits. The CLEM workflow we propose opens new avenues for the ultrastructural characterization of synapses with defined molecular signature characterizing their identity or activation profile in response to certain stimuli or behaviors. Another milestone to better model the biophysics of synaptic integration will be to combine EM and quantitative super-resolution LM to measure the density and nano-organization of molecular species (e.g. AMPARs, NMDARs, voltage-dependent calcium channels) in specific populations of synapses in intact brain circuits. Combining circuit and super-resolution approaches through CLEM will be critical to refine large-scale circuit models [74,133,150] (but see [32]) and bridge the gap between molecular, system and computational neurosciences.

## MATERIALS AND METHODS

### Animals and *in utero* cortical electroporation

All animals were handled according to French and EU regulations (APAFIS#1530-2015082611508691v3). *In utero* cortical electroporation was performed as described previously [151]. Briefly, pregnant Swiss female mice at E15.5 (Janvier Labs, France) were anesthetized with isoflurane (3.5% for induction, 2% during the surgery) and subcutaneously injected with 0.1 mg/kg of buprenorphine for analgesia. The uterine horns were exposed after laparotomy. Electroporation was performed using a square wave electroporator (ECM 830, BTX) and tweezer-type platinum disc electrodes (5mm-diameter, Sonidel). The electroporation settings were: 4 pulses of 40 V for 50 ms with 500 ms interval. Endotoxin-free DNA was injected using a glass pipette into one ventricle of the mouse embryos at the following concentrations: pH1SCV2 TdTomato: 0.5 μg/μL and pCAG EGFP-GPHN: 0.3 μg/μL. All constructs have been described before [98].

### Cortical slice preparation

Electroporated animals aged between postnatal day P84 and P129 were anesthetized with ketamin 100 mg/kg and xylazin 10 mg/kg, and intracardiacally perfused with first 0.1 mL of heparin (5000 U.I/mL, SANOFI), then an aqueous solution of 4% w/v paraformaldehyde (PFA) (Clinisciences) and 0.5% glutaraldehyde (GA) (Clinisciences) in 0.1 M phosphate-buffered saline (PBS). The fixative solution was made extemporaneously, and kept at ice-cold temperature throughout the perfusion. The perfusion was gravity-driven at a flow rate of about 0.2 ml/s, and the total perfused volume was about 100 ml per animal. Brains were collected and post-fixed overnight at 4°C in a 4% PFA solution. 30 μm-thick coronal brain sections were obtained using a vibrating microtome (Leica VT1200S).

### Fluorescence microscopy of fixed tissue

Slices containing electroporated neurons were trimmed to small (5-10 mm^2^) pieces centered on a relatively isolated fluorescent neuron, then mounted in a custom-made chamber on #1.5 glass coverslips. The mounting procedure consisted in enclosing the slices between the glass coverslip and the bottom of a cell culture insert (Falcon, ref. 353095) adapted to the flat surface with a silicon O-ring gasket (Leica) and fixed with fast-curing silicon glue (see panel A in S1 Fig). Volumes of GFP and tdTomato signals were acquired in 12 bits mode (1024×1024 pixels) with z-steps of 400 nm using an inverted Leica TCS SP8 confocal laser scanning microscope equipped with a tunable white laser and hybrid detectors and controlled by the LAF AS software. The objective lenses were a 10X PlanApo, NA 0.45 lens for identifying electroporated neurons and a 100X HC-PL APO, NA 1.44 CORR CS lens (Leica) for higher magnification images. GFP-GPHN puncta with a peak signal intensity at least four times above shot noise background levels were considered for CLEM.

### Placement of DAB fiducial landmarks

Following confocal imaging, slices were immersed in a solution of 1 mg/mL 3,3’-diaminobenzidine tetrahydrochloride (DAB, Sigma Aldrich) in Tris buffer (0.05 M, pH 7.4). The plugin “LAS X FRAP” (Leica) was used to focus the pulsed laser in the tissue in custom patterns of 10-to-20 points using 100% power in 4 wavelengths (470 to 494nm) for 30s-60s per point at 3 different depths: the top of the slice, the depth of the targeted soma, then the bottom of the slice (surface closest to the objective). DAB precipitates were imaged in transmitted light mode. Slices were subsequently rinsed twice in Tris buffer and prepared for electron microscopy.

### Tissue preparation for serial block-face scanning electron microscopy (SBEM)

Using a scalpel blade under a M165FC stereomicroscope (Leica), imaged tissue slices were cut to ~1mm^2^ asymmetrical pieces of tissue centered on the ROI, and then kept in plastic baskets (Leica) through the osmification and dehydration steps. Samples were treated using an osmium bridging technique adapted from the NCMIR protocol (OTO) [152]. The samples were washed 3 times in ddH_2_O and immersed for 1□hour in a reduced osmium solution containing 2% osmium tetroxide and 1.5% potassium ferrocyanide in ddH_2_O. Samples were then immersed for 20□minutes in a 1% thiocarbohydrazide (TCH) solution (Electron Microscopy Science) prepared in ddH_2_O at room temperature. The samples were then post-fixed with 2% OsO_4_ in ddH_2_O for 30□minutes at room temperature and colored *en bloc* with 1% aqueous uranyl acetate at 4□°C during 12□hours. Post-fixed samples were subjected to Walton’s *en bloc* lead aspartate staining at 60□°C for 30□minutes (Walton, 1979). After dehydration in graded concentrations of ice-cold ethanol solutions (20%, 50%, 70%, 90% and twice 100%, 5 minutes per step) the samples were rinsed twice for 10□minutes in ice-cold anhydrous acetone. Samples were then infiltrated at room temperature with graded concentrations of Durcupan (EMS) prepared without plastifier (components A, B, C only). In detail, blocks were infiltrated with 25% Durcupan for 30 minutes, 50% Durcupan for 30 minutes, 75% Durcupan for 2 hours, 100% Durcupan overnight, and 100% fresh Durcupan for 2 hours before being polymerized in a minimal amount of resin in a flat orientation in a sandwich of ACLAR^®^ 33C Films (EMS) at 60 °C for 48 hours. Samples were mounted on aluminum pins using conductive colloidal silver glue (EMS). Before curing, tissue blocks were pressed parallel to the pin surface using a modified glass knife with 0° clearance angle on an ultramicrotome (Ultracut UC7, Leica), in order to minimize the angular mismatch between LM and SEM imaging planes. Pins then cured overnight at 60°C. Samples were then trimmed around the ROI with the help of fluorescent overviews of the ROI within their asymmetrical shape. Minimal surfacing ensured that superficial DAB landmarks were detected at the SBEM before block-facing.

### SBEM acquisition

SBEM imaging was performed with a Teneo VS microscope (FEI) on the ImagoSeine imaging platform at Institut Jacques Monod, Paris. The software MAPS (Thermo Fisher Scientific) was used to acquire SEM images of targeted volumes at various magnifications. Acquisition parameters were: 1,7830 kV, 500 ns/px, 100 pA, 40 nm-thick sectioning and 8200×8200 pixels resolution with either 2.5 nm or 25 nm pixel size for high- and low-magnification images, respectively. Placing an electromagnetic trap above the diamond knife to catch discarded tissue sections during days-long imaging sessions was instrumental to achieve continuous 3DEM acquisitions.

### Image segmentation

Dendrites were segmented from SBEM stacks using the software Microscopy Image Browser (MIB) [153]. 3D reconstruction was performed with the software IMOD [154] (http://bio3d.colorado.edu/imod/). 3D spine models were imported in the software Blender (www.blender.org) for subsampling and the quantification of spine section areas along their main axis was done with in-house python scripts. Other measurements were performed using IMOD and in-house python scripts.

### Tissue preparation for tissue shrinkage estimation

Two female mice (21 days postnatal) were used for the analysis of tissue shrinkage induced by chemical fixation. Mice were decapitated and their brains were rapidly removed. The brains were transferred to an ice-cold dissection medium, containing (in mM): KCl, 2.5; NaHCO_3_, 25; NaH_2_PO_4_, 1; MgSO_4_, 8; glucose, 10, at pH 7.4. A mix of 95% O_2_ and 5% CO_2_ was bubbled through the medium for 30 min before use. 300-μm-thick coronal brain sections were obtained using a vibrating microtome (Leica VT1200S). Small fragments of the SSC were cut from those slices and fixed either by immersion in an ice-cold PBS solution containing 4% PFA and 0.5% GA, or in frozen with liquid nitrogen under a pressure of 2100 bars using a high pressure freezing system (HPM100, Leica). For HPF-frozen samples, the interval between removal of the brain and vitrification was about 7 min. Cryo-substitution and tissue embedding were performed in a Reichert AFS apparatus (Leica). Cryo-substitution was performed in acetone containing 0.1% tannic acid at −90°C for 4 days with one change of solution, then in acetone containing 2% osmium during the last 7h at −90°C. Samples were thawed slowly (5°C/h) to −20°C and maintained at −20°C for 16 additional hours, then thawed to 4°C (10°C/h). At 4°C the slices were immediately washed in pure acetone. Samples were rinsed several times in acetone, then warmed to room temperature and incubated in 50% acetone-50% araldite epoxy resin for 1h, followed by 10% acetone-90% araldite for 2h. Samples were then incubated twice in araldite for 2h before hardening at 60°C for 48h. As for chemically fixed sections, they were post-fixed for 30 min in ice-cold 2% osmium solution, rinsed in PBS buffer, dehydrated in graded ice-cold ethanol solutions and rinsed twice in ice-cold acetone, before undergoing the same resin infiltration and embedding steps as HPF-frozen samples. After embedding, ultrathin sections were cut in L2/3 of the SSC, orthogonally to the apical dendrites of pyramidal neurons, 200-300 μm from the pial surface using an ultramicrotome (Ultracut UC7, Leica). Ultra-thin (pale yellow) sections were collected on formwar-coated nickel slot grids, then counterstained with 5% uranyl acetate in 70% methanol for 10 min, washed in distilled water and air dried before observation on a Philips TECNAI 12 electron microscope (Thermo Fisher Scientific).

### Measurement of shrinkage correction factors

Ultra-thin sections of both HPF-frozen tissues and chemically-fixed tissues were observed using a Philips TECNAI 12 electron microscope (Thermo Fisher Scientific). Cellular compartments contacted by a pre-synaptic bouton containing synaptic vesicles and exhibiting a visible electron-dense PSD at the contact site, but no mitochondrion within their cytosol were identified as dendritic spine heads. Cross-section areas of random spine heads and the curvilinear lengths of their PSD were quantified in both conditions using the softwares MIB and IMOD. N = 277 spine head sections were segmented in HPF-frozen cortical slices from two female mice, and N = 371 spine head sections were segmented in chemically fixed cortical slices originating from the same two mice. Chi-square minimization was used between spine head cross-section area distributions in HPF or OTO conditions to compute average volume shrinkage and correction factors. PSD areas were not corrected as they exhibited no shrinkage.

### Computation of the diffusional neck resistance

The diffusional resistance of spine necks W_neck_ was measured as follows. Using IMOD, we first modeled in 3D the principal axis of each spine neck as an open contour of total length L_axis_ connecting the base of the neck to the base of the spine head. Using Blender, we interpolated each spine neck path linearly with 100 points. We named P(ℓ) the plane that bisected the spine neck model orthogonally to the path at the abscissa ℓ, and A(ℓ) the spine neck cross-section within P(ℓ). In spines containing a spine apparatus (SA), we corrected A(ℓ) by a scaling factor β(ℓ) = 1 – (D_SA_/D_spine_)^2^(ℓ), where D_SA_/D_spine_(ℓ) is the local ratio of SA and neck diameter. We measured D_SA_/D_spine_ orthogonally to the neck path in 10 SA+ spines and in three different locations per spine on SBEM images: at the spine stem ℓ/L_axis_ = 0.1), at the center of the spine neck (ℓ/L_axis_ = 0.5), and at the stem of the head (ℓ/L_axis_ = 0.9). D_SA_/D_spine_ was 44% ± 11%, 31% ± 8% and 37% ± 8% respectively, and fluctuations were not statistically significant. We then divided each SA+ spine neck in thirds and scaled their neck cross-section areas along neck axis A_SA+_(ℓ) = β(ℓ)A(ℓ) before computing W_neck_ = ∫ dℓ/A(ℓ) for all spines, using Simpson’s integration rule.

### Multi-compartment electrical model

All simulations were implemented in Python using NEURON libraries [155] and in-house scripts. Ordinary differential equations were solved with NEURON-default backward Euler method, with Δt = 0.05 ms. Scripts and model definition files are available in a GitHub repository: https://github.com/pabloserna/SpineModel. Biophysical constants were taken from the literature as follows: membrane capacitance C_m_ = 1μF/cm^2^ [38]; cytosolic resistivity ρ = 300 Ω.cm [85,156]; synaptic conductivities were modeled as bi-exponential functions g(t) = A g_max_ (e^−t/t_2_^ − e^−t/t_1_^) where A is a normalizing constant and (t_1_, t_2_) define the kinetics of the synapses: GABAergic conductance (t_1_, t_2_) = (0.5, 15) ms, AMPAR-dependent conductance (t_1_, t_2_) = (0.1, 1.8) ms, NMDAR-dependent conductance (t_1_, t_2_) = (0.5, 17.0) ms (ModelDB: https://senselab.med.yale.edu/ModelDB/). The magnesium block of NMDA receptors was modeled by a voltage-dependent factor [157]. Remaining free parameters comprised: the leaking conductivity g_m_ (or, equivalently, the membrane time constant T_m_); the peak synaptic conductance per area: g_AMPA_, g_NMDA_, g_GABA_; the total membrane area of the modeled neuron. These parameters were adjusted so that signal distributions fitted published electrophysiological recordings [110,113,116,158,159]. In more detail, we first set up one “ball-and-stick” model per segmented spine (N = 390). The dendrite hosting the modeled spine was generated as a tube of diameter d_dendrite_ = 0.87 μm, and length L_dendrite_ = 140 μm. This dendrite is split in three parts, the 2 μm-long middle one harboring the modeled spine. To account for the passive electrical effects of neighboring spines, the membrane surfaces of both the proximal and distal sections of the studied dendrite were scaled by a correction factor γ = 1 + <A_spine_> d_spine_ / πd_dendrite_ = 3.34, with the density d_spine_ = 1.63 spine.μm^−1^ and the average spine membrane area <As_pine_> = 3.89 μm^2^. We calibrated synaptic conductances type by type, by fitting the signals generated in the whole distribution of 390 models to published electrophysiological recordings. The AMPA conductances of all excitatory synapses were set proportional to ePSD area and scaled by the free parameter g_A_. In each model, we activated the AMPAR component of excitatory synapses and monitored the amplitude of resulting EPSCs in the soma. The average EPSC amplitude was adjusted to 58 pA [110,158], yielding a scaling factor g_A_ = 3.15 nS/μm^2^, which takes into account the average number of excitatory contacts per axon per PN in L2/3 of mouse SSC: N_ePSD/axon_ = 2.8 [110]. The average AMPA synaptic conductance was 0.42 nS. The leakage resistance was fitted to 65 MΩ [114], yielding a total membrane surface of the modeled neurons: A_mb,total_ = 18550 μm^2^. The NMDA conductances of all excitatory synapses were set proportional to ePSD area and scaled by the free parameter g_N_. In each model, we activated both NMDA and AMPA components of excitatory synapses and fitted the amplitude ratio between the average AMPA+NMDA and AMPA-only responses to 1.05 [158], yielding g_N_ = 3.4 nS/μm^2^. The GABA conductances of all inhibitory synapses were set proportional to iPSD area and scaled by the free parameter g_G_. In this case, we set the neuron to a holding potential of 0 mV and the reversal potential of chloride ions (E_Cl_^−^) to −80 mV. Then, we activated shaft inhibitory synapses and monitored the amplitude of resulting IPSCs in the soma. The amplitude of the average GABAergic conductance was set to 1 nS [38,114,115], yielding a scaling factor g_G_ = 5.9 nS/μm^2^, which takes into account the average number of inhibitory contacts per axon per PN in L2/3 of mouse SSC: N_iPSD/axon_ = 6 [114]. Considering inhibition, since E_Cl_^−^ is regulated on timescales exceeding 100 ms [160] and we modeled signals in the 10 ms timescale, we could set E_Cl_^−^ as a constant parameter of our steady state model. Calcium influxes were modeled in spines as a result of the opening of NMDARs and voltage-dependent calcium channels (VDCCs). Since we simulated signals that remained below the threshold for eliciting dendritic spikes [35,74], we did not include VDCCs in dendrites, and monitored calcium transients exclusively in spine heads. The dynamics of L-, N- and Q-type VDCCs were obtained from ModelDB (accession n°: 151458), and their conductivities were scaled to the head membrane area of each spine, A_head_, excluding synaptic area(s). VDCC-type ratios and calcium conductivities were adjusted by fitting the average amplitude of calcium concentration transients to 20% of the NMDA conductance [161]. Calcium uptake from cytosolic buffers was set to 95% to yield an average amplitude of Ca^2+^ concentration transients of 0.7 μM [162].

### Bootstrapping

To simulate a large number of spine-spine interactions with limited redundancy, our distribution of spines was expanded using a “smooth” bootstrapping method [121]. Specifically, the dataset (i.e. a matrix of dimensions N x N_f_) was re-sampled to generate a new matrix of dimension M x N_f_, where N is the number of spines, N_f_ is the number of selected features, and M is the final number of synthetic spines. M rows were randomly selected in the original dataset and zero-centered, feature-dependent Gaussian noise was added to each element of the matrix (excluding absolute quantities, e.g. number of PSDs or presence of SA). To determine appropriate noise amplitude for each parameter, a synthetic set of M=500 spines was generated from the original dataset, including Gaussian noise with an arbitrary amplitude σ, on one selected parameter. This new feature distribution was compared to the original distribution using a 2-sample Kolmogorov-Smirnov test (KS-test), and this procedure was repeated 1000 times for each set value of σ. A conservative noise level (σ = 10%) was sufficient to smear parameter distributions while the fraction of synthetic sets that were statistically different from the original set (p < 0.05, KS-test) remained 0 over 1000 iterations. σ = 10% was valid for all relevant features, and we assumed that such a small noise amplitude would minimally interfere with non-linear correlations in our dataset. Synthetically generated spines were then used to simulate elementary synaptic signaling using in-house python scripts. We also used bootstrapping to estimate standard deviations in our simulations.

### Statistics

No statistical methods were used to predetermine sample size. We used a one-way ANOVA on our 4 datasets (S1 Table) to test that inter-neuron and inter-mice variability were small enough to pool all datasets together (S2 Table). We used Kolmogorov-Smirnov test to determine that all measured morphological parameters followed a log-normal distribution (S2 Table). We used Mann–Whitney U test for statistical analyses of morphological parameters, except when comparing the probability for SiSs and DiSs to harbor SA, for which we used Pearson’s χ^2^ test. All results in the text are mean ± SD. In Fig 6,Fig 7 and Fig 9, we plot medians as solid lines, as they better describe where log-normal distributions peak. Shaded areas represent 68% confidence intervals, which span approximately one standard deviation on each side of the mean.

## ACKNOWLEDGEMENTS

We thank members of the Triller and Charrier laboratories (IBENS, Paris, France), Vincent Hakim and Boris Barbour for insightful discussions. This work was supported by INSERM, the Agence Nationale de la Recherche (ANR-13-PDOC-0003 and ANR-17-ERC3-0009 to C.C.), the European Research Council (ERC starting grant 803704 to C.C.), the Labex Memolife (901/IBENS/LD09 to O.G.), the company NIKON France via the CIFRE program (convention n° 2015/1049 to O.G.) and the European Union’s Horizon 2020 Framework Programme for Research and Innovation under the Specific Grant Agreement No. 785907 (Human Brain Project SGA2 to P.S.). We acknowledge the ImagoSeine imaging facility (Jean-Marc Verbavatz and Rémi Leborgne, Jacques Monod Institute, Paris, France) for SBEM availability, service and support. We thank Liesbeth Hekkings (ThermoFisher Scientific) for technical support. We are grateful to the IBENS Imaging Facility (France BioImaging, supported by ANR-10-INBS-04, ANR-10-LABX-54 MEMO LIFE, and ANR-11-IDEX-000-02 PSL* Research University, ‘‘Investments for the future”; NERF 2011-45; FRM DGE 20111123023; and FRC Rotary International France).

## AUTHOR CONTRIBUTIONS

O.G. developed the 3D-CLEM workflow, performed the experiments, segmented images and analyzed data. P.S. coded the model and analyzed data. N.A. and M.F. carried out the *in utero* electroporations. P.R. trained O.G. in EM and carried out part of the TEM imaging. O.G., P.R., A.T. and C.C. designed the study. O.G., P.S. and C.C. interpreted the results, prepared the figures, and wrote the manuscript. A.T. and C.C. provided funding.

