## Supporting information for "Morphologically constrained modeling of spinous inhibition in the somato-sensory cortex"

### FiGURES

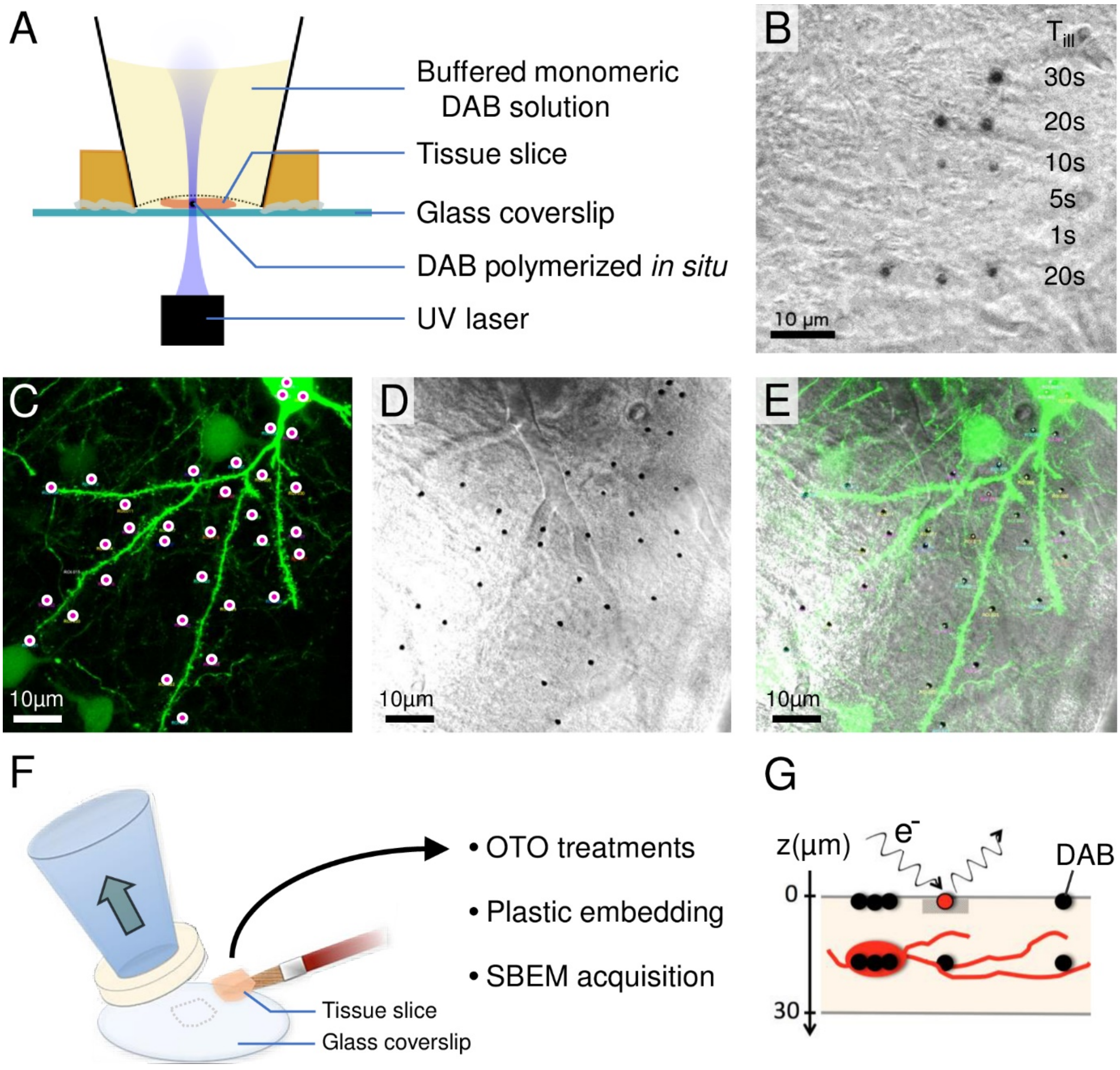


**S1 Fig. ROI landmarking strategy for 3D-CLEM using DAB and a detachable chamber.**

(A) Schematic of ROI landmarking using DAB photo-oxidation. The tissue slice is held against a glass coverslip in a solution of DAB using a detachable chamber. Confocal imaging and ROI landmarking are performed within the same microscopy setup.

(B) Transmitted light image of a cortical slice labelled with DAB precipitates. Varying the duration of UV illumination ($T_{ill}$) allows adjusting DAB spot size.

(C) Example of labeling pattern around an optically-isolated fluorescent neuron. DAB was photo-precipitated by focusing UV light for 40s at each highlighted location (pink dots in yellow circles).

(D) Transmitted light image of the same field of view after DAB photo-precipitation.

(E) Overlay showing DAB precipitates in D (dark spots) arranged similarly to the UV focusing pattern in C.

(F) Schematic of slice retrieval after ROI landmarking. Detaching the chamber (wide arrow) allows taking the sample to EM preparation steps, i.e. Osmium-TCH-Osmium post-fixation (Deerinck et al., 2010), dehydration, resin infiltration and plastic embedding.

(G) ROI recovery within the SBEM. DAB precipitates (circles) generated at the surface of the sample mark the (x,y) coordinates of the ROI. They are detected with an electron beam (e-) before block-facing the sample, and acquiring SBEM images in targeted volumes. The DAB pattern generated at the depth of the targeted cell (in red) allows identifying it retrospectively.

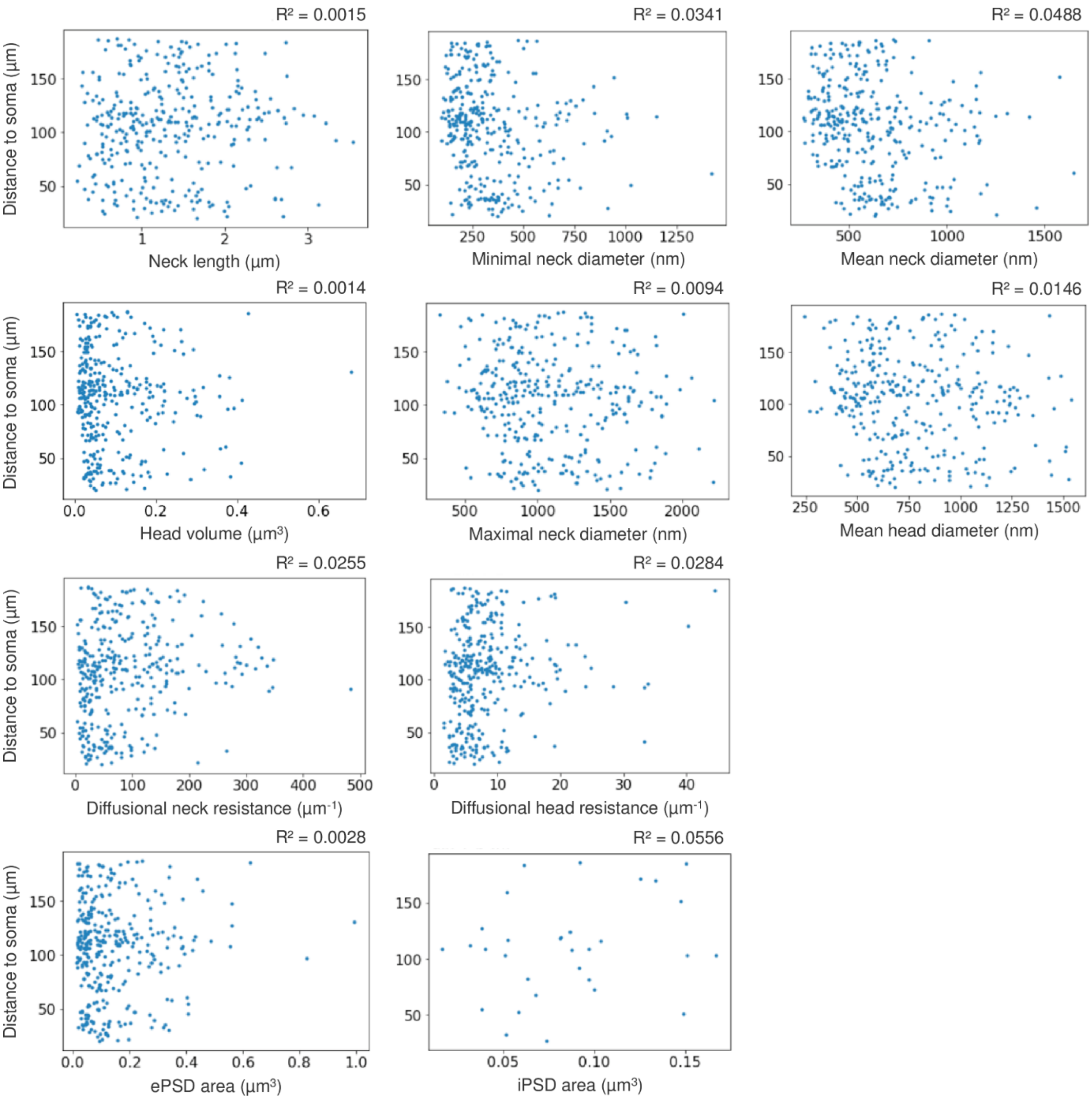


**S2 Fig. Distance between spine and soma does not correlate with other measured parameters.**

Distance between spine and soma as a function of all measured morphological parameters, for all segmented spines. No parameter exhibit a linear correlation with the distance between spine and soma.

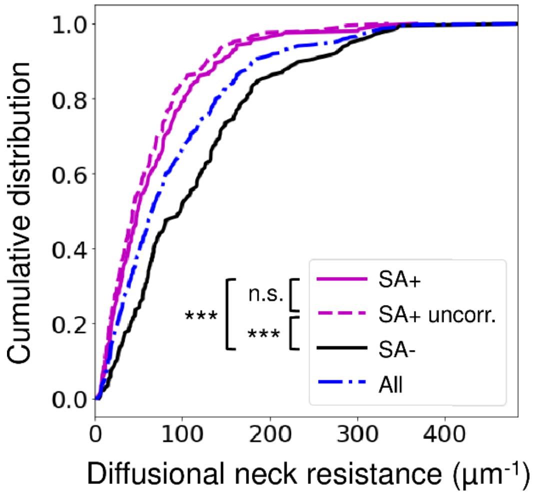


**S3 Fig. Effect of the spine apparatus on diffusional neck resistance.**

Distribution of the diffusional neck resistance ($W_{neck}$) for spines devoid of apparatus (SA-) or containing a spine apparatus (SA+) calculated using neck morphology. SA+ uncorr.": SA was not taken into account for the calculation of $W_{neck}$for these SA+ spines (p=0.1 compared to corrected $W_{neck}$). ***p < 0.001 calculated using Mann-Whitney test.

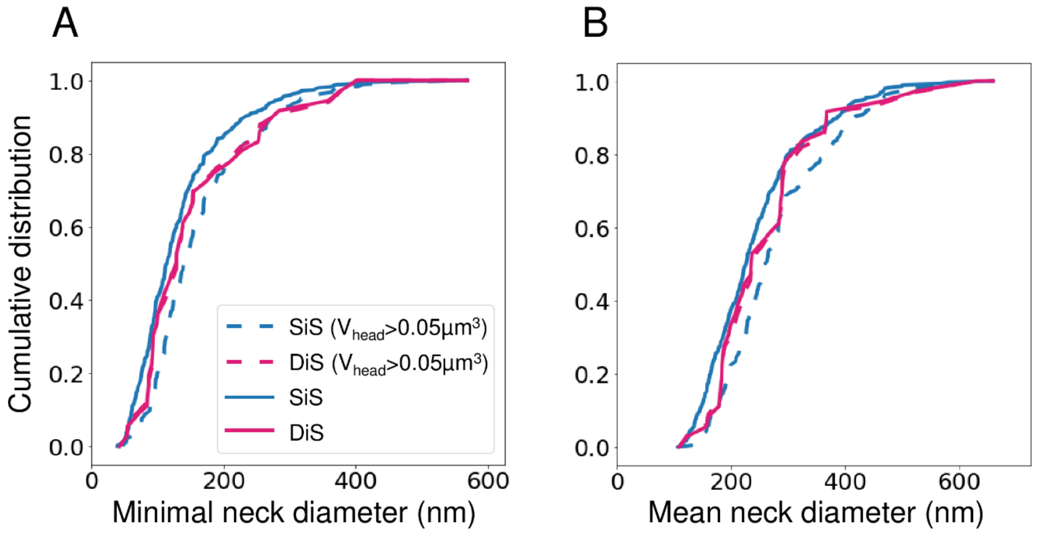


**S4 Fig. Comparison of SiS and DiS neck diameter.**

Quantification of minimal neck diameter (A) and mean neck diameter (B) for all spines (N=349 SiSs and 37 DiSs; solid lines) and for spines with $V_{head}$ > 0.05 µm$^3$ (N=186 SiSs and 34 DiSs; dashed lines). p > 0.05 calculated using Mann-Whitney tests.

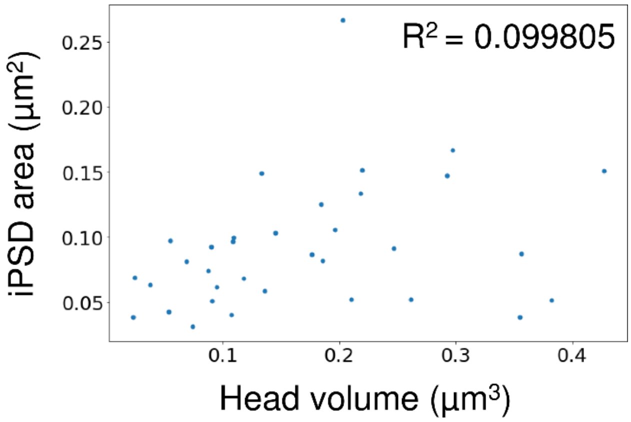


**S5 Fig. Inhibitory PSD area as a function of DiS head volume.**

Inhibitory PSD area as a function of spine head volume for N=37 DiSs. Linear regression: $R^2 < 0.1$ .

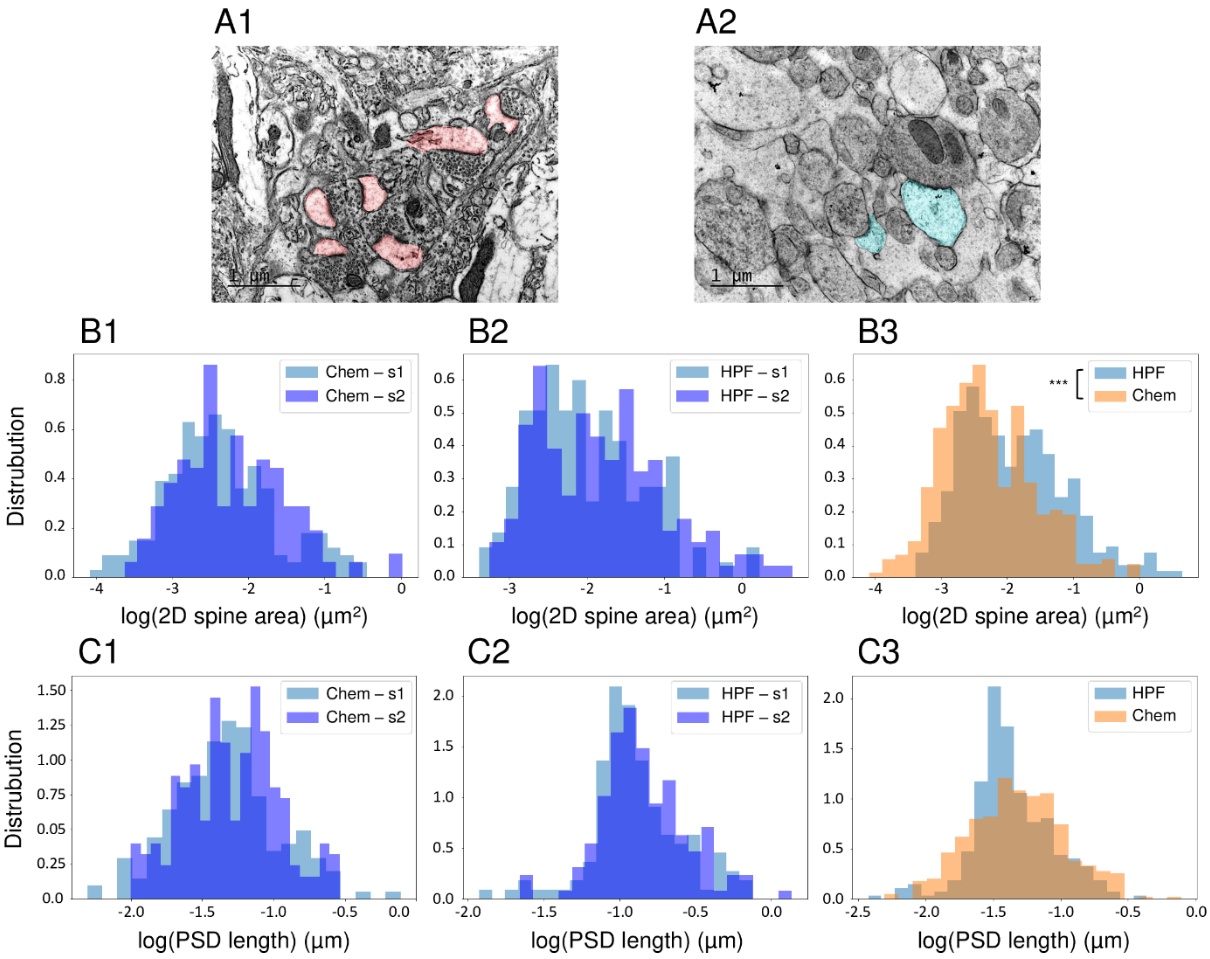


**S6 Fig. Fixation-induced shrinkage of spine heads but not PSD area.**

(A) TEM images of L2/3 SSC acute slices from the same N=2 young adult mice (s1 and s2; 21d postnatal), upon either chemical fixation with aldehydes (A1), or physical fixation with high-pressure freezing (HPF) (A2). Spine head section areas (indicated in red in A1 and in light blue in A2) and lengths of PSDs were segmented for quantification. Scale bars: 1 µm.

(B) Normalized histograms of spine head cross-section areas for chemically-fixed (Chem) samples (N = 194 for s1 and 178 for s2; $p=0.13$) or HPF samples (N = 128 for s1 and 150 for s2; $p=0.052$) in B1 and B2 respectively. Data from s1 and s2 displayed no statistical difference and were pooled together in B3 to compare area distribution between HPF (blue) and chemically-fixed tissue (orange). Head cross-section areas were 36% smaller in chemically fixed samples (orange) than in HPF samples (blue), implying ~49% head volume shrinkage ($p < 10^{-8}$). (C) Normalized histograms of PSD section lengths for chemically-fixed ($p=0.44$) and

HPF samples (p=0.17) in C1 and C2 respectively. PSDs were not significantly deformed by chemical fixation (C3; p=0.10). Only significant ($p < 0.05$) p-values are shown. ***$p < 0.001$ calculated using Mann-Whitney test.

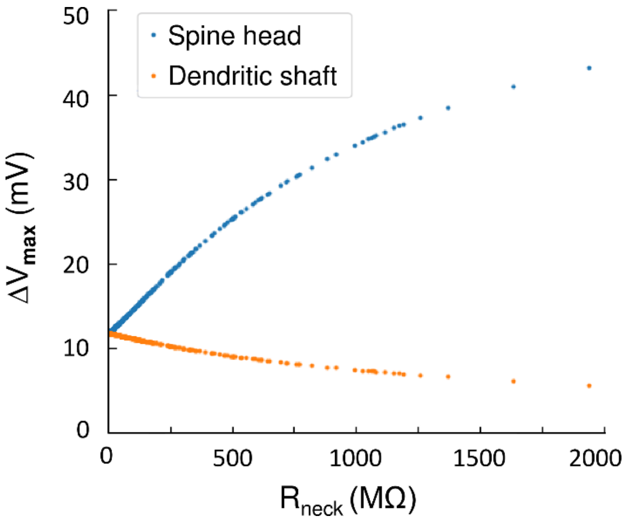


**S7 Fig. Influence of $R_{neck}$ on $\Delta V_{max}$ in the spine head and dendritic shaft.**

Plot of the EPSP amplitude $\Delta V_{max}$ elicited in a dendritic spine while varying its neck resistance ($R_{neck}$) and keeping all other parameters constant. Increasing $R_{neck}$ causes $\Delta V_{max}$ to increase in the spine head (blue) and to decrease in the dendritic shaft (orange).

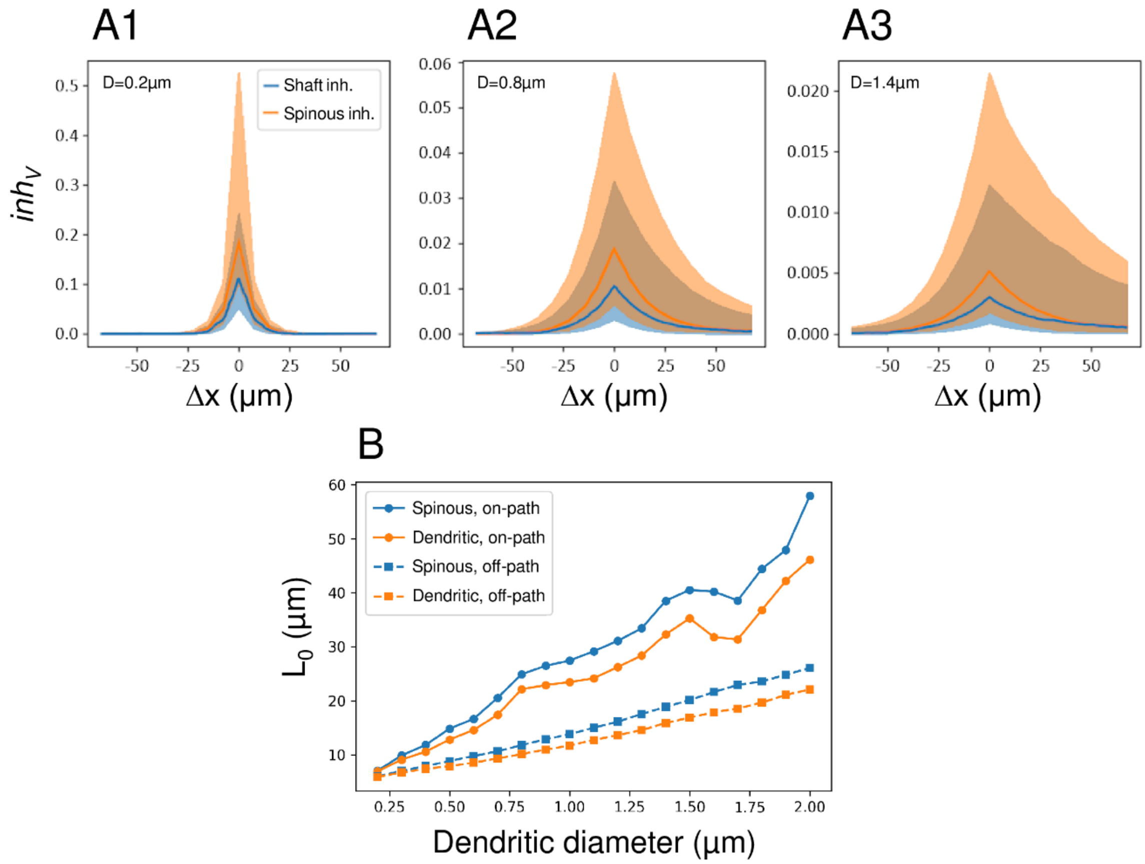


**S8 Fig. Impact of dendritic diameter on electrotonic lengths.**

(A) Voltage inhibition $inh_V$ in the spine as a function of $\Delta$x, for different values of the dendritic diameter (D): A1: D = 0.2 µm; A2: D = 0.8 µm; A3: D = 1.4 µm. Solid lines represent medians and shaded areas represent 68% confidence intervals.

(B) Decaying length $L_0$ of the exponential fits of the "on-path" and "off-path" parts of $inh_V$ as a function of the dendritic diameter, where "on-path" and "off-path" correspond to the positive and negative range of $\Delta$x, respectively.

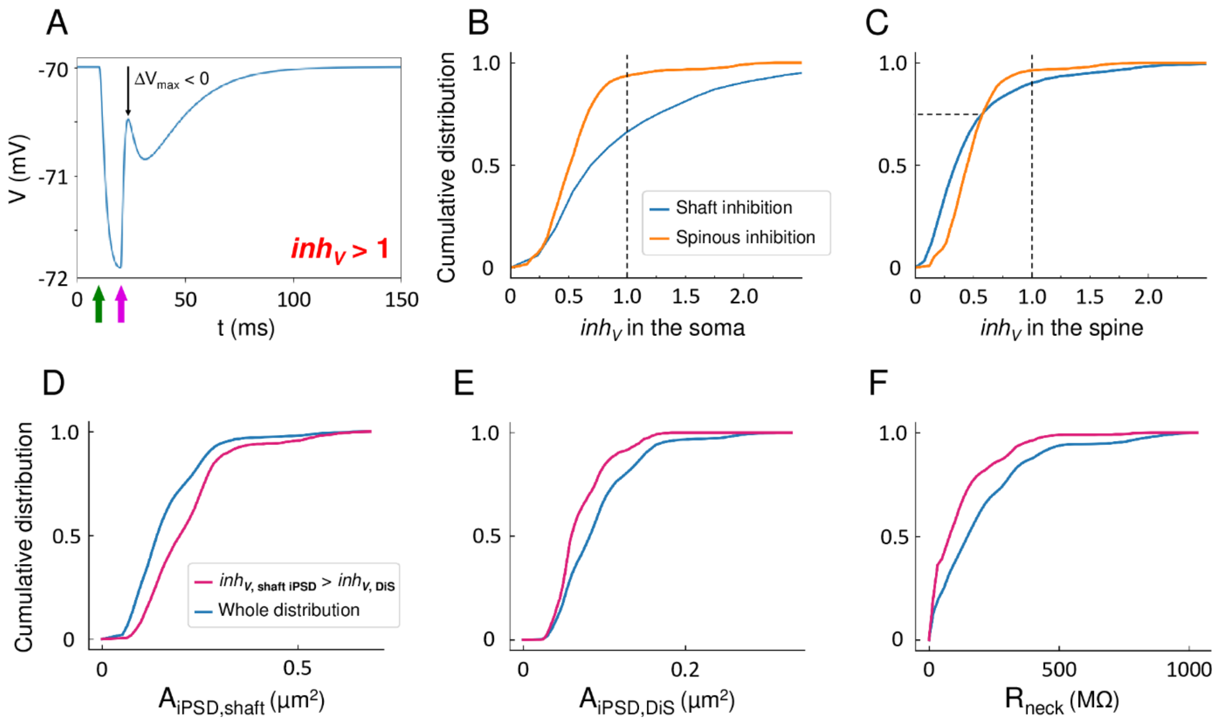


**S9 Fig. Impact of hyperpolarizing inhibition on EPSP integration.**

(A) Example time-course of a hyperpolarizing IPSP summed with a weak EPSP ($inh_V > 1$ and $\Delta V_{max} < 0$).The IPSP (green arrow) was elicited 10ms after the EPSP (magenta arrow) to clarify the contribution of each PSP.

(B-C) Estimation of $inh_V$ in the soma (B) or in the spine head (C) (N=3700 iterations of the model). The IPSP was elicited either in a DiS (orange) or in the dendritic shaft (blue).

(D-F) Distribution of shaft iPSD area (D), DiS iPSD area (E) and spine neck resistance (F) in the N=3700 simulations. Blue: distributions for all iterations. Magenta: distributions for the iterations in which dendritic shaft inhibition yielded a higher value of $inh_V$ in the spine than spinous inhibition.

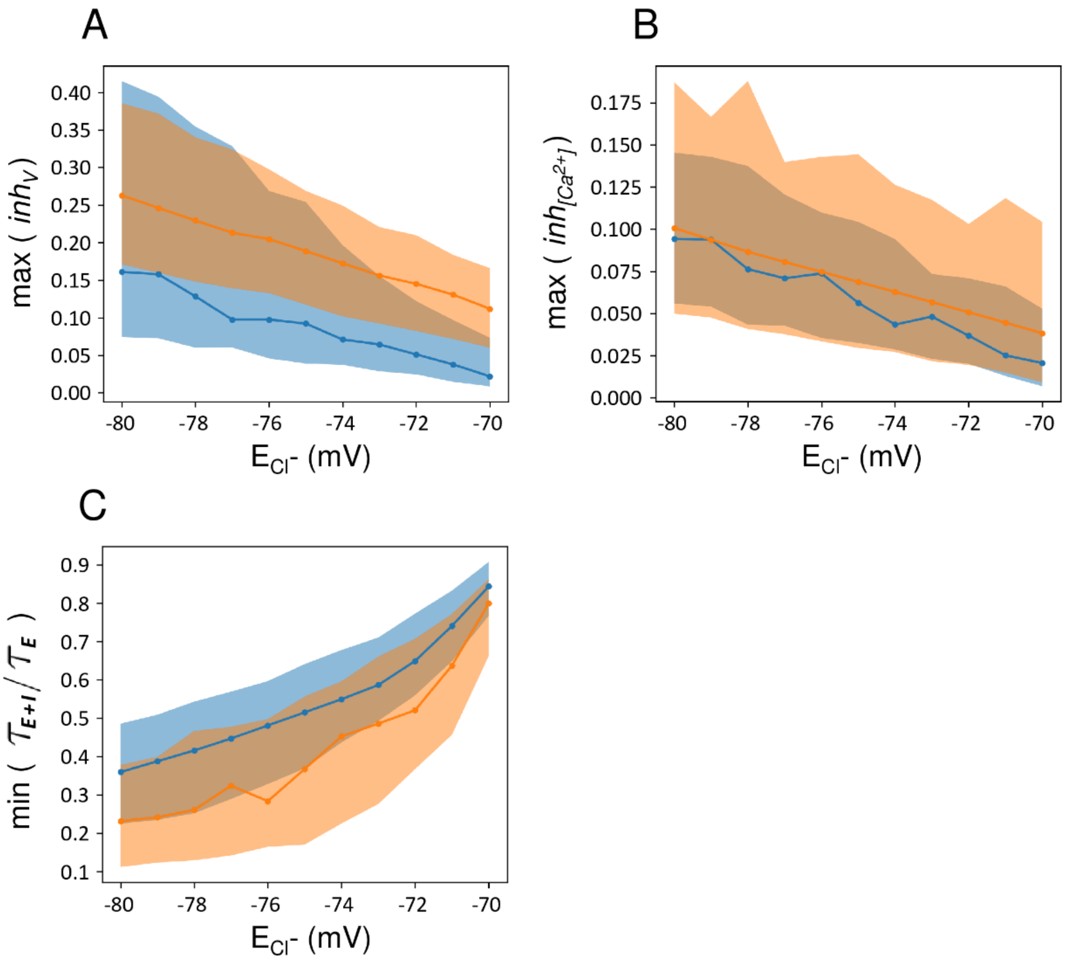


**S10 Fig. Impact of chloride reversal potential on EPSP inhibition.**

Estimation of maximal voltage inhibition (A), maximal calcium inhibition (B), and the minimal ratio of depolarization decay time in the spine with or without inhibition, $\tau_{E+I}/\tau_E$ (C) as a function of the chloride reversal potential $E_{Cl^-}$ and depending on the placement of the iPSD either on the dendritic shaft (blue) or on the spine (orange). Solid lines represent medians. Shaded areas represent 68% confidence intervals.

### TABLES

| Dataset | Mouse ID | Sex | Age (days) | Number of segmented dendrites | Number of segmented spines |
|---|---|---|---|---|---|
| **180609_133** | 1 | F | 84 | 1 | 48 |
| **180618_222** | 2 | M | 84 | 2 | 170 |
| **180626_232** | 2 | M | 84 | 3 | 131 |
| **180723_322** | 3 | F | 129 | 2 | 41 |

**S1 Table. Sample description.**

| Feature | p-value for KS-test against log-normal distribution | p-value for ANOVA across datasets on log(feature) |
|---|---|---|
| **Head volume** | 0.2526 | 0.39454 |
| **Neck length** | 0.2562 | 0.0189006 |
| **ePDS area** | 0.1679 | 0.191623 |
| **iPSD area** | 0.9967 | 0.420379 |
| **Minimal neck diameter** | 0.295 | 0.0115228 |
| **Mean neck diameter** | 0.7657 | 0.109704 |
| **Neck resistance** | 0.2002 | 0.0101641 |
| **Head length** | 0.1912 | 0.412945 |
| **Maximal head diameter** | 0.4483 | 0.00806033 |
| **Mean head diameter** | 0.4721 | 0.0131462 |
| **Head surface** | 0.664 | 0.617778 |

**S2 Table. Log-normality and inter-sample variability tests.**

KS-test: Kolmogorov-Smirnov test.

| Output | Contribution to the variance of the output | | | | | |
|---|---|---|---|---|---|---|
| | in SiSs | | | in DiSs | | |
| $\Delta V_{max,\ soma}$ | $A_{ePSD}$ 0.8907 | $L_{dend}$ 0.0641 | $R_{neck}$ 0.0005 | $A_{ePSD}$ 0.7668 | $L_{dend}$ 0.1468 | $R_{neck}$ 0.0092 |
| $\Delta V_{max,\ spine\ head}$ | $A_{ePSD}$ 0.602 | $R_{neck}$ 0.1882 | $L_{dend}$ 0.0004 | $A_{ePSD}$ 0.4683 | $R_{neck}$ 0.3787 | $L_{dend}$ 0.0095 |
| $\Delta V_{max,\ shaft} / \Delta V_{max,\ spine\ head}$ | $R_{neck}$ 0.9687 | $L_{dend}$ 0.008 | $A_{ePSD}$ 0.0043 | $R_{neck}$ 0.9682 | $L_{dend}$ 0.0128 | $A_{ePSD}$ 0.0038 |
| $\Delta[Ca^{2+}]_{max,\ spine\ head}$ | $A_{ePSD}$ 0.3023 | $R_{neck}$ 0.0878 | $L_{dend}$ 0 | $A_{ePSD}$ 0.4514 | $R_{neck}$ 0.0941 | $L_{dend}$ 0.0002 |

**S3 Table. Contribution of morphological parameters to the variance of $\Delta V_{max}$ and $\Delta[Ca^{2+}]_{max}$.**

Highest-ranking input parameters ($A_{ePSD}$, $R_{neck}$ and $L_{dend}$) are sorted by decreasing contribution to the variance of the simulation outputs, as estimated with a GLM. Numbers indicate to which proportion input variables accounted for the variance of considered output.

| Output | Contribution to the variance of the output | | | | | | | |
|---|---|---|---|---|---|---|---|---|
| | $E_{Cl^-}$ = -70mV | | | | $E_{Cl^-}$ = -80mV | | | |
| min($\tau_{E+I} / \tau_E$) | $A_{iPSD}$ 0.6414 | $L_{dend}$ 0.2208 | $R_{neck}$ 0.0300 | $A_{ePSD}$ 0.0176 | $A_{iPSD}$ 0.4453 | $L_{dend}$ 0.1726 | $R_{neck}$ 0.1049 | $A_{ePSD}$ 0.0730 |
| max($inh_V$) | $A_{iPSD}$ 0.3990 | $R_{neck}$ 0.3683 | $L_{dend}$ 0.0014 | $A_{ePSD}$ 0.0001 | $A_{ePSD}$ 0.2434 | $A_{iPSD}$ 0.2135 | $R_{neck}$ 0.0087 | $L_{dend}$ 0.0016 |
| max($inh_{[Ca^{2+}]}$) | $A_{ePSD}$ 0.2738 | $R_{neck}$ 0.2236 | $A_{iPSD}$ 0.0187 | $L_{dend}$ 0.0085 | $A_{ePSD}$ 0.2429 | $R_{neck}$ 0.2046 | $A_{iPSD}$ 0.1213 | $L_{dend}$ 0.0222 |

**S4 Table. Contribution of morphological parameters to the variance of summed signals.**

Highest-ranking input parameters ($A_{ePSD}$, $A_{iPSD}$, $R_{neck}$ & $L_{dend}$) are sorted by decreasing contribution to the variance of the simulation outputs as estimated with a GLM. Numbers indicate to which proportion input variables accounted for the variance of considered output.

### DATA

**S1 Data. Quantification of the morphology of spines and synapses.** Legends are included in the rightmost columns of each sheet of the .xlsx file.
